# Disproportion among reticulon-like 16 (RTNLB16) splice variants disrupts growth and decreases sensitivity to ABA and senescence in Arabidopsis

**DOI:** 10.1101/2023.12.18.572161

**Authors:** Tami Khazma, Dikla Levi, Hiba Waldman Ben-Asher, Gad Miller

## Abstract

The Reticulon family proteins (RTNs) are membrane-spanning proteins found in the endoplasmic reticulum (ER) with diverse functions, such as ER membrane morphogenesis, vesicle formation, and trafficking. The plant-specific reticulon-like protein family (RTNLBs) comprises multiple members, yet their functions remain poorly understood. The Arabidopsis RTNLB16 gene has seven splice variants, each encoding seven distinct protein isoforms.

We identified an Arabidopsis mutant (Salk_122275/*rtnlb16-1*) as a knockout for the upper coding frame, isoform 7, of RTNLB16 while overexpressing the other six isoforms through the CaMV 35S promoter at the left border of the T-DNA insertion. *rtnlb16-1* exhibits distinctive growth retardation and reduced chlorophyll levels. Under photoperiodic long day (16:8 h) conditions, activation of the 35S promoter intensifies *RTNLB16* expression in the mutant, resulting in profound growth inhibition. Conversely, growth under continuous low-light (CLL) conditions restrains the overexpression and significantly mitigates *rtnlb16-1* phenotype. Confocal microscopy experiments revealed the localization of RTNLB16:GFP in the tubular ER network, plasmodesmata, and potentially in Golgi bodies.

Peculiarly, RTLB16/*rtnlb16* heterozygote plants exhibit non-Mendelian reduced fertility, suggesting potential involvement of RTNLB16 in reproductive development. Transcriptomics comparisons between *rtnlb16-1* and the wild type under CLL and 16:8h conditions revealed differential gene expression involved in salicylic acid, jasmonic acid, and abscisic acid responses, indicating activation of defense and osmotic stress responses contributing to the growth inhibition in the mutant. We further demonstrate that *rtnlb16* has decreased sensitivity to abscisic acid and enhanced tolerance to darkness-induced senescence.

Our findings highlight the importance of balanced expression among RTNLB16 isoforms for normal cellular and physiological activities in Arabidopsis. Additionally, our study underscores the significance of employing T-DNA mutants to investigate genes with multiple splice variants.

## Introduction

Reticulon (RTN) family proteins constitute a class of highly conserved endoplasmic reticulum (ER)-resident proteins recognized for their diverse functions in cellular processes. The family comprises reticulons (RTNs) in vertebrates and reticulon-like family proteins (RTNLs) in non-chordates and other Eukaryotes, including the plant subfamily of RTNLBs (Oertle et al., 2003). RTNs are integral membrane proteins featuring conserved reticulon homology domains (RHDs), a 200-amino-acid residue region at the C-terminus responsible for associating with the ER membrane. The RHD is composed of two hydrophobic regions, each 28-36 amino acids long, flanking a 60-70 residue hydrophilic loop (Yang and Strittmatter, 2007).

The continuous membrane system of the ER includes the nuclear envelope, flat sheets (cisternae) often studded with ribosomes, and a polygonal network of highly curved tubules extended throughout the cell (Goyal and Blackstone, 2013). The primary function of RTNs is their involvement in ER shaping, facilitating membrane curvature, and maintenance. Reticulons contribute to the formation of tubules and sheets within the ER, thereby influencing its overall morphology (Voeltz et al., 2006). RTNs exhibit various possible topologies in membranes, with the hydrophobic regions of the RHD spanning the membrane completely, partially, or looping within the membrane (Yang and Strittmatter, 2007). The hydrophobic portion of the RHD occupying the outer layer of the phospholipid bilayer can induce membrane curvature via hydrophobic wedging. As oligomers, RTNs bend the membrane by forming immobile arc-like scaffolds (Shibata et al., 2008; Goyal and Blackstone, 2013; Wang and Rapoport, 2019). In contrast to the highly conserved C-terminus, the N-terminal regions are highly variable, exhibiting little or no sequence homology, and are likely to interact with distinct proteins, conferring specific biological functions (Di Sano et al., 2012).

RTNs have been demonstrated to have diverse functions, participating in various cellular processes and responses. Reticulons are primarily implicated in regulating membrane dynamics and trafficking processes from the ER to the Golgi apparatus. They participate in vesicle formation, membrane fusion, and apoptosis (Yang and Strittmatter, 2007; Wang and Rapoport, 2019; Voeltz et al., 2006). RTN3 functions as a specific ER-phagy receptor responsible for the initiation and degradation of ER tubules (Grumati et al., 2017). Some RTNs are also found to interact with the plasma membrane. In the mammalian central nervous system (CNS), RTN4a/NogoA, localized in the plasma membrane, acts at the cell surface to inhibit neurite growth (Oertle et al., 2003; Sui et al., 2015).

The plant RTN gene family, RTNLBs, is larger and more diverse than in animals and yeasts. The RTNLB family includes many paralogues in the same plant species as a result of gene duplication (Oertle et al., 2003; Nziengui and Schoefs, 2009). Plant RTNLBs have been clustered into four phylogenetic groups that differ in the length of the variable C- and/or N-terminus (Nziengui and Schoefs, 2009). The Arabidopsis genome includes 21 RHD-containing RTNLB proteins with a similar structural organization to mammalian RTNs, suggesting similar functions. AtRTNLB2 and 4 were shown to associate with the ER similarly to animal RTNs but with different distribution patterns through the ER continuum. RTNLB2 is specifically located in the tubular sub-compartment of the ER, and RTNLB4 is also located in the lamellar cisternae (Nziengui et al., 2007).

Several Arabidopsis RTNLBs were identified as localized in ER-derived desmotubules lining the plasmodesmata pore, suggesting they play a role in their formation (Know 2015 – 26084919). Nearly all RTN/RTNL genes contain multiple introns and exons, and most are alternatively spliced into multiple isoforms (Oertle et al., 2003; Yang and Strittmatter, 2007). According to the TAIR database, the majority of the AtRTNLB genes have two or more splice variants predicted to produce different protein isoforms, suggesting distinct functions. RTNLB11 and RTNLB16 have 7 splice variants, potentially producing 3 and 7 distinctive protein isoforms, respectively.

In this work, we identified the Salk_122277 T-DNA insertion mutant as having a unique disruption in the expression of *RTNLB16*, leading to severe growth retardation and a decreased level of chlorophyll. While in the wild type, all seven splice variants seem to be expressed, the *rtnlb16-1* is knocked out in isoform 7 while overexpressing the other 6 isoforms downstream through the CaMV 35S promoter located at the left border of the T-DNA. Under continuous light conditions that suppress the activity of the CaMV 35S promoter, the growth inhibition in *rtnlb16-1* is partially rescued. We show that mere overexpression of RTNLB16 does not recapitulate the growth phenotype on rtnlb16-1. Confocal microscopy revealed the cellular localization of RTNLB16:GFP in ER tubules, plasmodesmata, and potentially Golgi bodies. RNAseq analysis comparisons revealed the potential involvement of plant biotic and abiotic stress responses associated with salicylic acid (SA), jasmonic acid, (JA), and abscisic acid (ABA) signaling. We further showed that *rtnlb16-1* seed germination is insensitive to ABA, and the mutant is dark-stress tolerant.

Additionally, a previous study has wrongfully identified heterozygous Salk_122277 as a mutant impaired in the essential mitochondrial manganese superoxide dismutase gene (MSD1) deficient in embryo sac development and fertilization (Martin et al., 2013). This work also rectifies that mistake.

## Results

Our initial interest in studying mitochondrial manganese SOD (MSD1) led us to investigate the Salk_122275 T-DNA line. According to the SIGnAL T-DNA express mapping tool (http://signal.salk.edu/cgi-bin/tdnaexpress), this line contains a T-DNA insertion within the first exon of the MSD1 gene, disrupting its ORF. Previously dubbed *oiwa-1* mutant, Salk_12275, was previously reported to exhibit female gametophyte defects resulting in seed abortion in the heterozygote (Martin et al., 2013). This outcome aligns with expectations for essential genes like MSD1. Surprisingly, we isolated homozygous salk_122275 plants displaying retarded growth, indicating that the disruption in salk_122275 is not lethal (Figure 1b). Immunoblot analysis with anti-MSD1 antibodies and SOD enzymatic activity assays indicated wildtype-like MSD1 expression and activity in Salk_122275 homozygote plants (Fig. S1). These results proved that Salk_122275 is not a SOD1 mutant as previously claimed by Martin et al. 2003.

**Fig 1.**
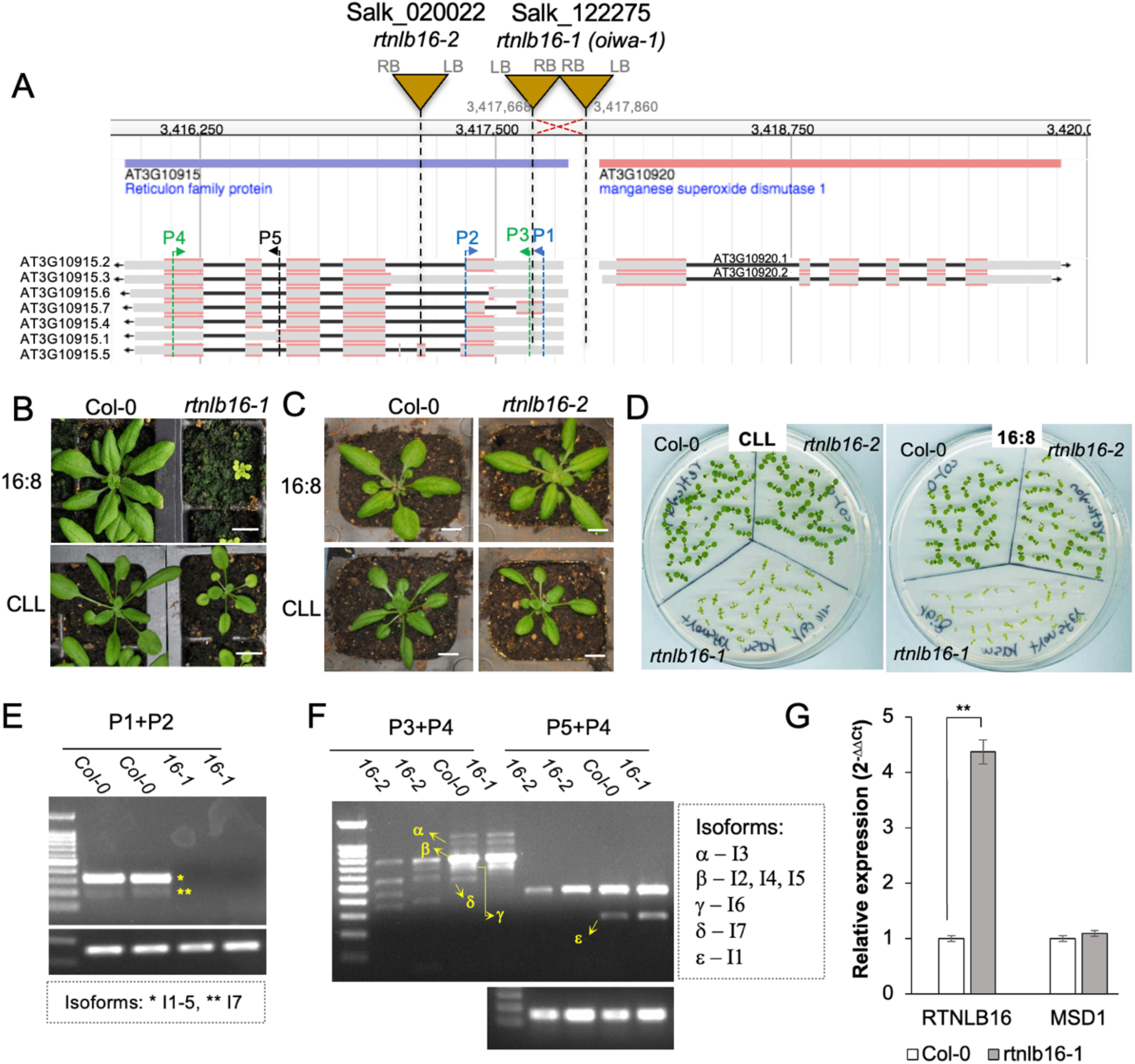
Genotype and phenotype of rtnlb16 T_DNA insert mutants. A. A genome browser map of RTNLB16 and MSD1 genes on chromosome 3. The gene model splice variants presented below show exons (red), UTRs (grey), and introns (black). Triangles indicate the T-DNA inserts with a vertical dashed line pointing to the position of integration in the gene models. Primers (P1-P5) used for the PCR assays and their orientation is indicated with arrows. B and C, Four-week-old WT and mutant plants germinated and continued growing on soil under the indicated light regime. D. Seven days old seedlings growing on solid MS media under both light conditions. E and F, Semi-qRT-PCR results of RTNLB16 expression profile, upper gels. Actin 2 expression, depicted below, was used for evaluating cDNA levels. Identified amplicons of specific splice variants are indicated with asterisks or Greek letters. G. qRT-PCR results comparing expression in leaves of CLL-grown plants.

PCR genotyping of Salk_122275 with the left border primer (LBb1) and the insertion site flanking primers produced two left border amplicons, indicating that the T-DNA was integrated in a double concatenated form with a head-to-head configuration (RB-RB) (Fig. 1a, Fig. S2A,B). Sequencing these two PCR products allowed us to accurately map the insertion site of the double inverted T-DNA in Salk_122275 to the RTNLB16 gene, the neighboring gene to MSD1, which is inversely located (head-to-head) on the opposite strand (Fig. 1a, Fig. S3). Integration of the inverted T-DNA double insertion led to a 191 nucleotides deletion, trimming off 120 bp from the upstream of RTNLB16 and 71 bp from the short intergenic region shared with MSD1 that spans the promoter of both genes (Fig. 1a, Fig. S3). The deletion within the RTNLB16 sequence trimmed part of the 5’UTR of isoforms 1-7, and the first 35 nucleotides in the upstream CDS of isoform 7 (Fig. 1A, Fig. S3). Under long-day conditions (16:8 hours light:dark, 80-100µE), Salk_122275/*rtnlb16-1* exhibits severe growth retardation and pale green leaves, which are significantly alleviated under continuous low light intensity (24 hours, 40µE (CLL)) conditions (Fig. 1B, D).

We isolated an additional mutant, *rtnlb16-2*, with a T-DNA insertion in the first intron common to isoforms 1, 2, 4, 6, 7, the second exon of isoform 5, and the 5’UTR of isoform 3 (Fig. 1A, Fig. S1C). However, *rtnlb16-2* displayed a growth phenotype like the wild type under 16:8 and CLL conditions (Fig. 1C, D).

Semi-quantitative RT-PCRs prepared from leaves of CLL-grown plants revealed that *rtnlb16-1* lacks expression of isoform 7. Yet, cDNA amplicons obtained with a primer set covering most of the gene indicated that isoforms 1-6 are expressed in this mutant (Figure 1E, F). This finding immediately suggested that the expression of six *RTNLB16* isoforms is driven by the CaMV-35S promoter located close to the left border of the pROK2 DNA vector (Ulker et al., 2008). Quantitative RT-PCR in plants grown under CLL revealed a 4-fold increase in *RTNLB16* transcript levels in *rtnlb16-1* compared with the WT. In contrast, the level of *MSD1* remained unchanged and equal to the WT (Fig. 1G). Semi-quantitative RT-PCR revealed that *rtnlb16-2* lacks the expression of isoforms 1, 3, and 5, and shows a reduced level of isoform 6 (Fig. 1F). Taken together, the results indicate that *rtnlb16-1* is not a simple loss-of-function null allele but rather suggests its complex phenotype is the result of a combination of the loss of isoform 7 and constitutive expression of the other isoforms.

We further assessed the differences in chlorophyll levels under the two light conditions using the ‘delayed fluorescence’ whole-plant live imaging protocol as a proxy (Gould et al., 2009; see materials and methods). Under 16:8 conditions, the chlorophyll level in *rtnlb16-1* was approximately half that of the wild type. Under CLL conditions, the chlorophyll level in the mutant increased by 18%, while in the wild type, it decreased by 17% compared with 16:8 conditions (Fig. 2). These results suggest that, similar to the growth inhibition phenotype, the chlorophyll content in rtnlb16-1 is dependent on photoperiodism.

**Fig 2.**
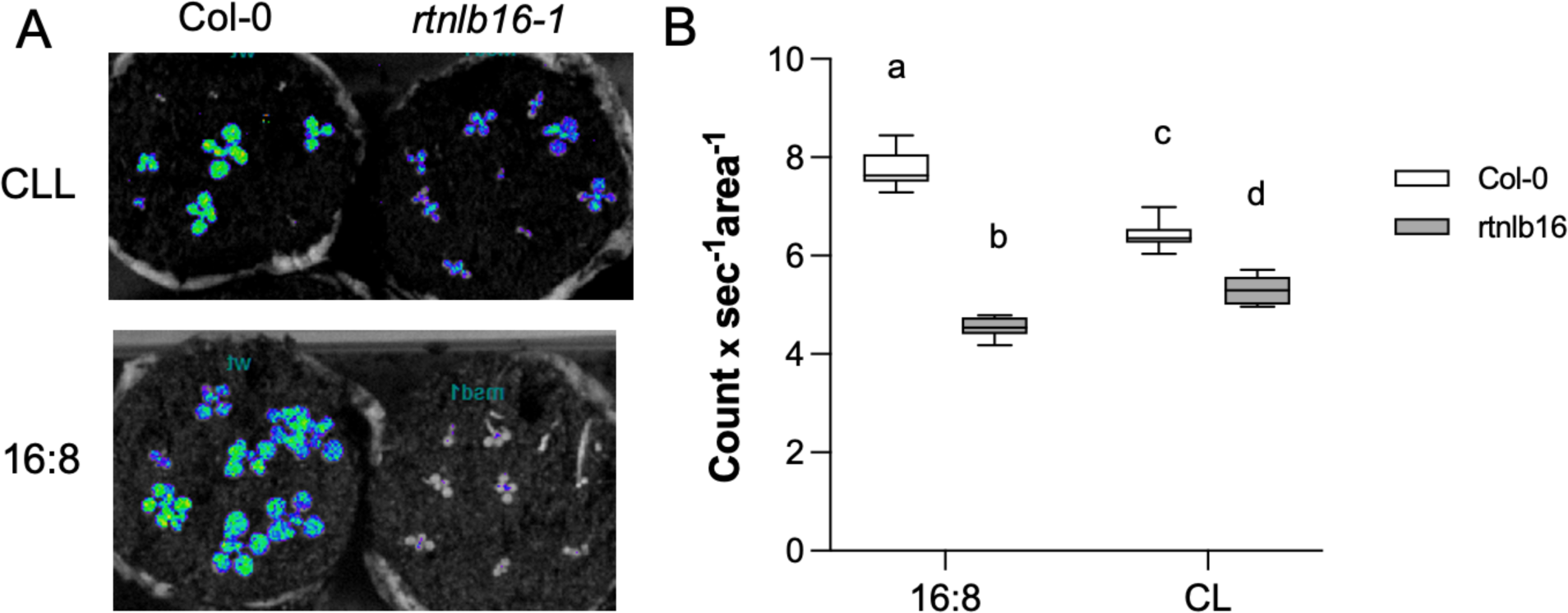
Chlorophyll evaluation in 2-weeks-old rosettes using delayed fluorescence imaging with the NightShade imager. A. Representing photos. B. Quantification of photons emitted from excited chlorophyll. Each sample average consisted of 6 replicates. The letters above boxes indicate statistical significance at the level of P<0.01 analyzed by two-way ANOVA and Tukey’s multiple comparisons test.

Heterozygous Salk_122275 shows no apparent defects during the vegetative growth phase and is indistinguishable from the wild type (Fig. S4), indicating that *rtnlb16-1* is a recessive allele. Since (Martin et al., 2013) reported a 50% abortion rate in the heterozygous, we reevaluated this phenotype by scoring seed set averages in plants grown under CLL conditions. The seed set in heterozygote plants exhibited relatively large variation with an average of 27 seeds/silique, 35% lower than in the wild type, which averaged 42 seeds (Fig. 3). Yet, the increased abortion rate in the heterozygotes, evident as undeveloped seeds (Fig. 3a), is significantly lower than the normal Mendelian 50% expected in cases of essential genes. In contrast with heterozygote *rtnlb16-1*, the seed set in the homozygous rtnlb16-1 was closer to the wild type with 37 seeds/silique (Fig. 3b). These results indicate non-Mendelian inheritance for this phenotype, suggesting that the increase in ovule abortion or failure in fertilization was not directly caused by the absence of isoform 7.

**Fig. 3.**
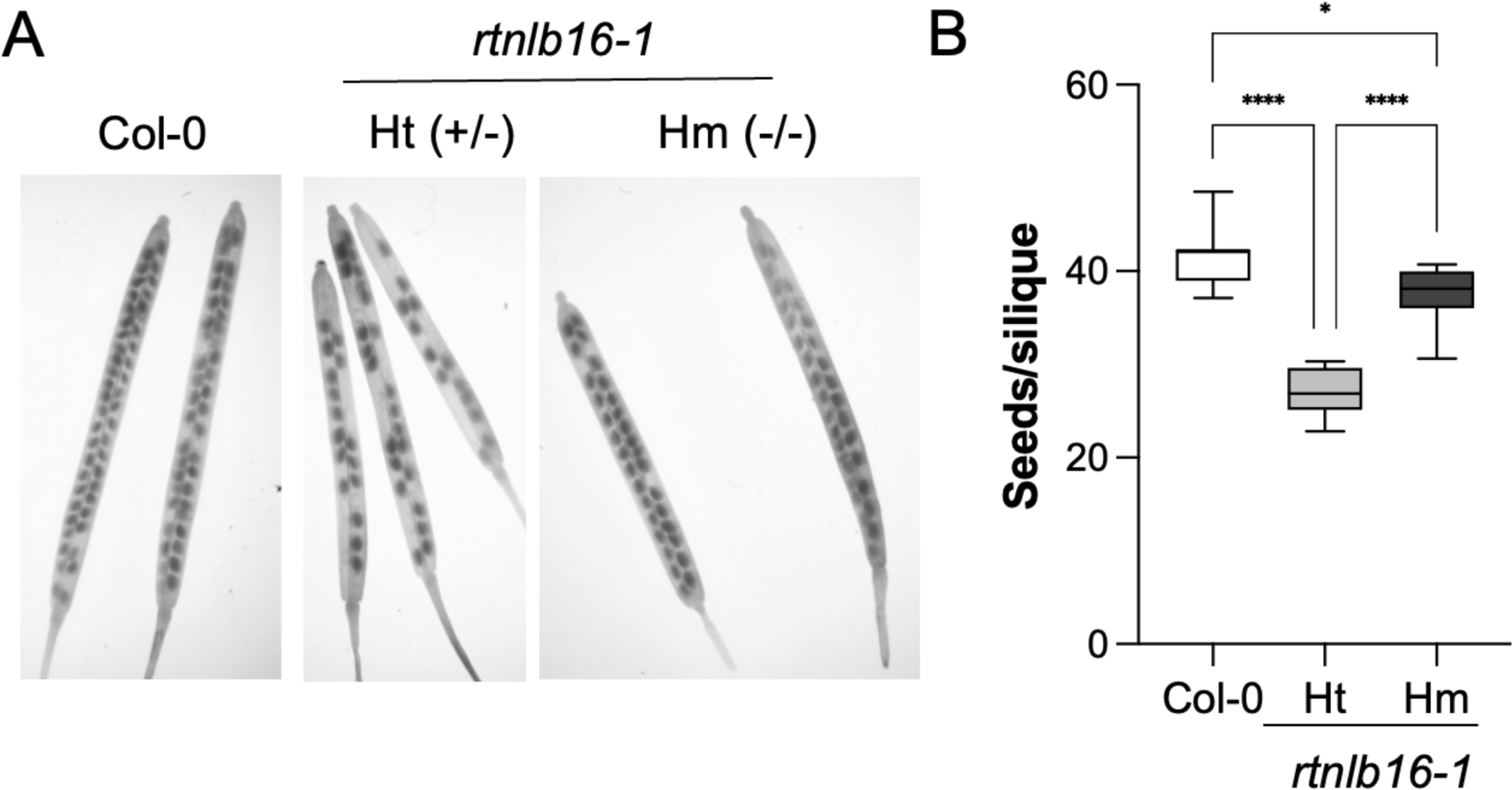
Reproductive phenotype of decreased seed set in *rtnlb16-1/oiwa-1*. A. Photos show siliques of six-week-old plants growing under CLL conditions. B. Box plot graph. Mean values and errors represent the average from 100 siliques (10 siliques from 10 plants). * and *** indicate P<0.05 and P<0.001 in one-way ANOVA and Tukey’s multiple comparisons test.

To gain a deeper understanding of the function and complexity of RTNLB16, isoform 5 protein, the primary splice variant product, fused to GFP to determine its cellular localization. Transient co-expression assays of RTNLB16.5:GFP with an endoplasmic reticulum mCherry marker (ER-rk; Nelson et al., 2007) in N. benthamiana leaves epidermal cells, revealed a partial overlap between the two fluorescent proteins. Confocal microscope cross-section images in leaf epidermal cells indicated that RTNLB16 is localized in the ER tubular network (Fig. 4A-C, Fig. 5S). Yet, additional cytoplasmic localization of RTNLB16:GFP not overlapping with the ER mCherry appeared as ER-independent puncta (Fig. 4C, Fig. S5), potentially indicative of Golgi vesicles/bodies (Jung et al., 2012).

**Fig. 4.**
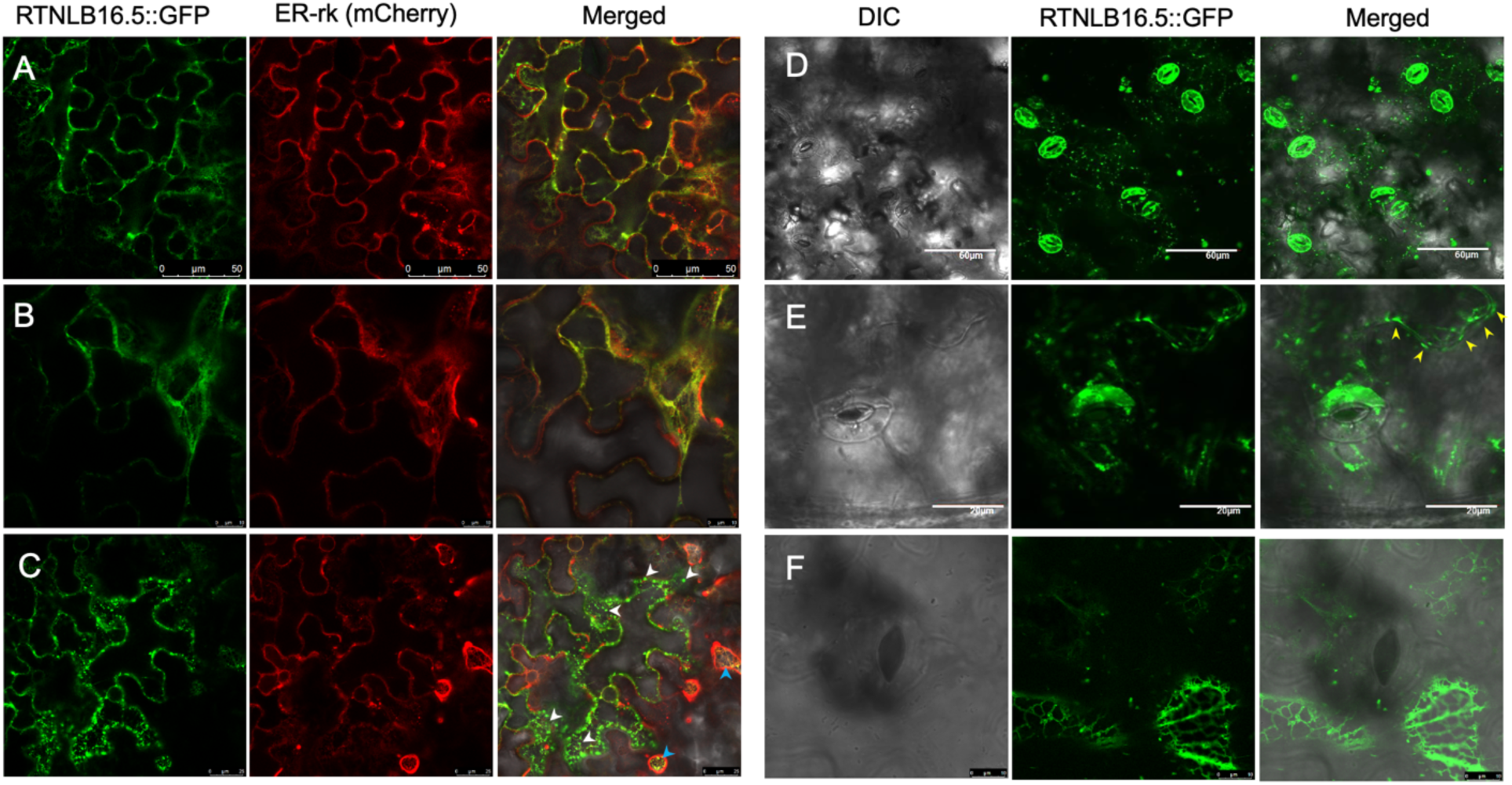
Cellular localization of RTNLB16.5:GFP. Confocal images of RTNLB16.5:GFP in N. benthamiana leaves trasiently co-expressed with an ER mCherry marker (A-C), and in trasgenic Arabidopsis (D-F). Blue arrows in C point to regions of complete overlap between the two fluorescent markers, and white arrows point to GFP puncta suspected as golgi bodies.

We further generated transgenic Arabidopsis plants expressing RTNLB16.5:GFP. In the leaf epidermis, the GFP signal was most prominent in guard cells, displaying an evident ER-network shape within the cytoplasm, suggesting that RTNLB16 function might be important for stomata activity (Fig. 4D,E). In pavement epidermal cells, the RTNLB16:GFP subcellular pattern revealed the distinct ER network layout of tubules and cisternae (Fig. 4F). Additionally, GFP-rich structures closely associated with the plasma membrane, resembling plasmodesma, were noticeable (PD; Fig. 4E, F). These results suggest a complex subcellular localization pattern for RTNLB16.

We then set to gain a better understanding of the relationship between the growth inhibition phenotype in *rtnlb16-1* and photoperiodism. To this end, plants germinated and initially grown under continuous low light (CLL) conditions were transferred to continue growing under 16:8 conditions until maturity at one- or two-week intervals. The size of the plants at 5 weeks of age was compared between transfer intervals, and plants were kept constantly at CLL. Results revealed a positive correlation between the size of the *rtnlb16-1* plant and the duration of time growing under the 16:8 photoperiodic growth condition (Fig. 5A). The diameter of mutant plants constantly growing under CLL conditions was 30% larger than plants of the same age following one week under 16:8, suggesting a rapid and drastic inhibition of growth upon exposure to photoperiodism. Moreover, the growth inhibition upon transfer to 16:8 and the release of inhibition when returning rtnlb16-1 plants to CLL were apparent within a day (Fig. S6).

**Fig. 5.**
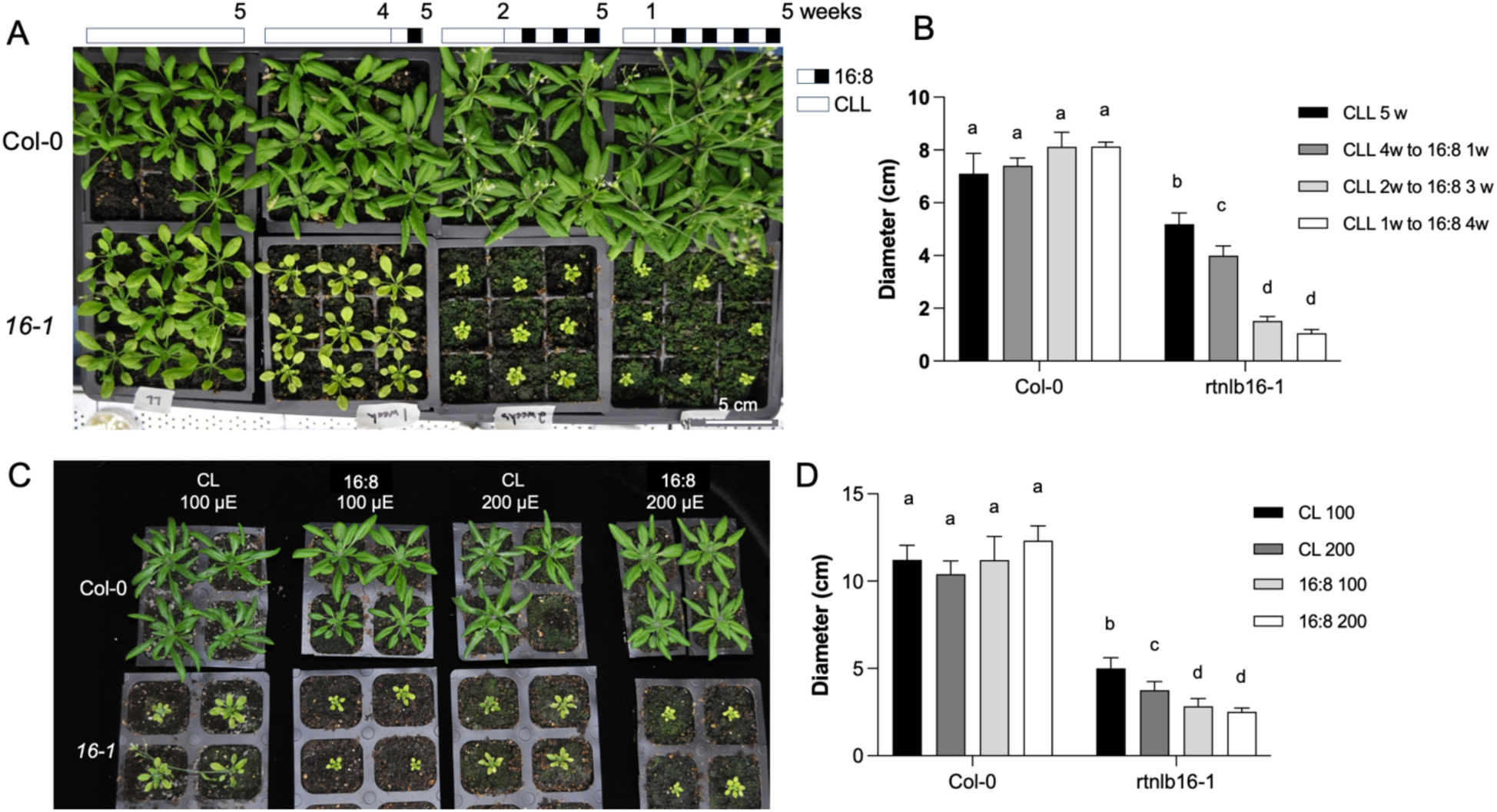
Light-dependent phenotype of rtnlb16-1. A and B, Relationship between time growing under CLL or 16:8 conditions: Plants germinated at CLL were transferred for increased duration of 1, 3, or 4 weeks to 16:8. A. Photo of plants side by side at 5 weeks old age. C and D, an experiment comparing the impact of light intensity and light regime on rtnlb16-1 phenotype at 4 weeks of age: Two weeks old plants grown under CLL were transferred to continuous light (CL) or 16:8 conditions with moderate light intensity of either 100 or 200 µE (µmol*S^-1^*m^-2^). Means and error bars of rosette diameter represented in B and D are an average of 8 replicates. The letters above the bars indicate statistical significance at the level of P<0.01 analyzed by two-way ANOVA and Tukey’s multiple comparisons test.

Because the intensity at 16:8 was nearly double that of CLL, we investigated whether the growth retardation of rtnlb16-1 is influenced by light intensity. Seedlings germinated at CLL conditions were subjected to 100 µE and 200 µE light intensities under 16:8 or continuous light (CL) conditions for five weeks. While the wild type did not display significant differences in size among the different conditions, *rtnlb16-1* plants were the smallest under 16:8, with no significant difference between the two light intensities. Under 100 µE CL, the diameter of rtnlb16-1 was 33% larger than under 200 µE and less than half the size of the wild type (Fig. 5B, C, Fig. S7). Similarly, the inflorescences of rtnlb16-1 were shorter under 16:8 than under CL, with photoperiodism having a greater effect than light intensity (Fig. S7). These results suggest the light-to-dark transition has the most critical effect on *rtnlb16-1* growth inhibition.

Interestingly, the CaMV 35S promoter, responsible for the high expression of *RTNLB16* isoforms in *rtnlb16-1* (Fig. 1G), is photoperiod-responsive, exhibiting weak expression in darkness and heightened expression during daylight (Schnurr and Guerra, 2000; Saidi et al., 2009). Therefore, we assessed the activity of the 35S promoter under continuous low light (CLL) and 16:8 conditions using transgenic Arabidopsis seedlings expressing GFP. GFP fluorescence imaging revealed that under 16:8, the GFP signal intensity was nearly twice that observed under CLL conditions (Fig. 6A). Subsequently, using semi-quantitative RT-PCR, we investigated whether the expression level of *RTNLB16* transcripts in the mutant differed under the two light regimes. Three-week-old plants grown under CLL were introduced to 16:8 for 1 and 3 days (Fig. S6A). Under both conditions, a higher-size PCR product corresponding to isoform 3 or the inclusion of intron 1 of isoforms 1-6 was detected in the mutant but not in the wild type (Figure 6B). On the third day following transfer to 16:8, the expression of all isoforms significantly increased in rtnlb16-1 compared to low expression in the wild type (Figure 6B). Interestingly, one day after the transfer to 16:8 (following the first nighttime period), the *RTNLB16* level in the wild type transiently increased, suggesting that the RTNLB16 promoter is also influenced by photoperiodism.

**Fig. 6.**
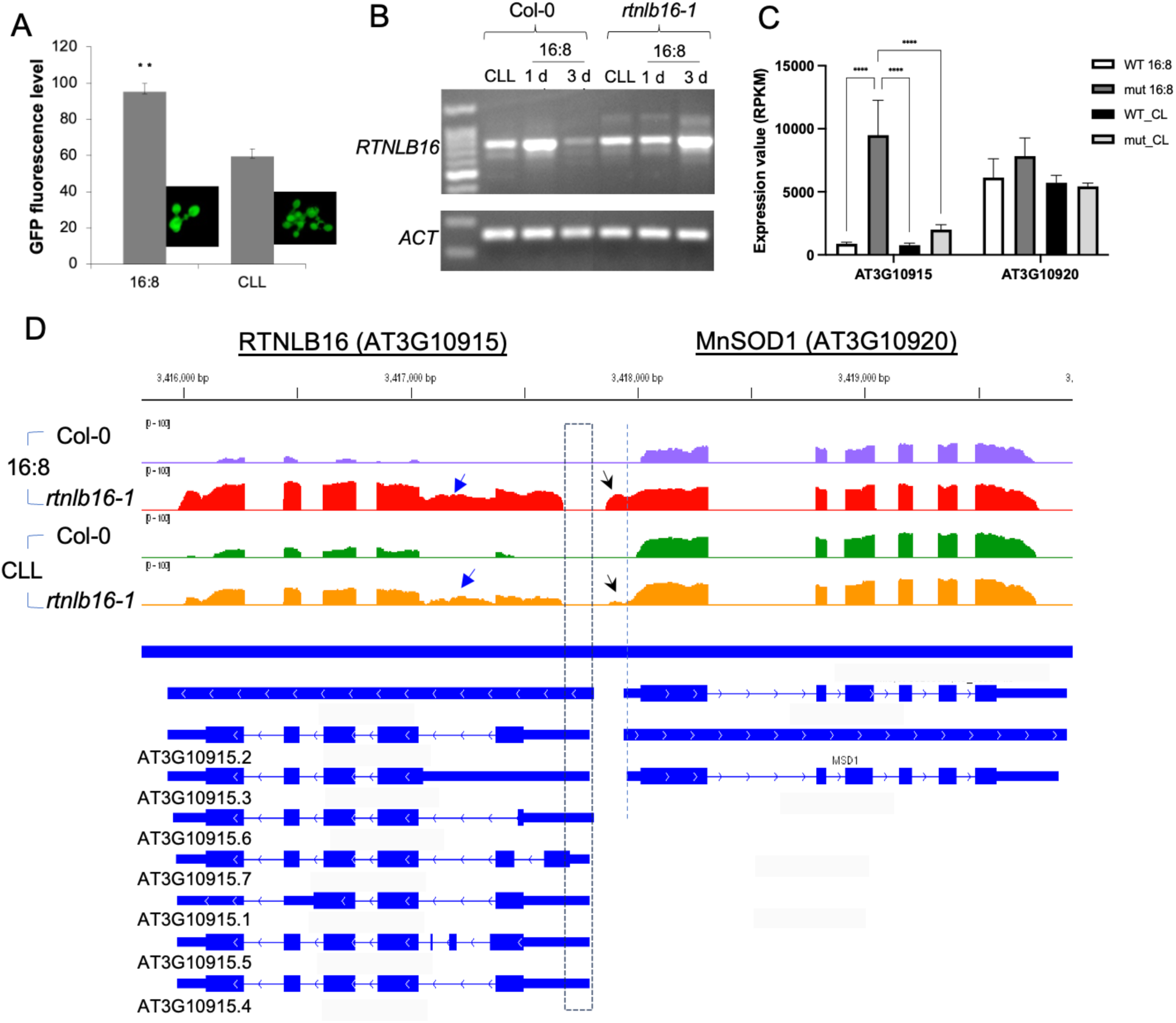
Increased expression of RTNLB16 in rtnlb16-1 is controlled by 35S promoter in a light-dependent mode. A. Live imaging quantification of GFP fluorescence level driven in two weeks old 35S:GFP transgenic plants grown under CLL or 16:8 conditions. Error bars represent the average of six replicates. ** indicate p value<0.01 in a student T-test. The representing GFP image is shown next to the respective histogram. B. Semin quantitative PCR results of RTNLB16 transcripts feom plants grown at CLL and after 1 and 3 days following transfer to 16:8 regime. C. Expression level of RTNLB16 and MSD1 in WT and rtnlb16-1 obtained from RNAseq analysis in CLL or 16:8 grown leaves. Normalized values are in Reads Per Kilobase per Million mapped reads (RPKM). Two-way ANOVA Tukey’s multiple comparisons test was conducted for each gene’s expression. **** indicate statistical significance with P<0.001. D. RNAseq reads coverage for RTNLB16 and MSD1 shown by Integrative Genomics Viewer (IGV). Blue arrows point to increased read coverage corresponding to transcripts of isoform 3 or the inclusion of intron 1 of isoforms 1, 2, 4, and 6. Black arrows point to reads coverage corresponding to miss-expression of MSD1 upstream promoter region likely due to activation by 35S promoter in the T-DNA insert.

To identify the transcriptional program responsible for the *rtnlb16-1* phenotype, we conducted an RNA sequencing experiment in 4-week-old plants grown under both light regimes. The *RTNLB16* transcript levels in the RNAseq data were 2.6- and 11-fold higher in the mutant under CLL and 16:8, respectively, compared to the wild type (Fig. 6 C). Integrated Genomic Viewer (IGV) display of the base coverage for RTNLB16 showed a distribution of sequencing reads that is enhanced in the mutant, covering all the exons apart from the segment overlapping with the upstream of isoform 7, which was deleted due to the T-DNA insertion (Fig. 6D). In addition, reads covering the intergenic promoter sequence of MSD1 exclusively generated in the mutant suggest activation by the 35S promoter at the left border on the MSD1 side. Yet, the level of *MSD1* in the mutant remained largely unchanged (Fig. 6C, D). These results confirm that the 35S promoter at the T-DNA insert left border controls the activation of *RTNLB16* in *rtnlb16-1* in a photoperiodic-dependent manner. Additionally, the results suggest a direct causal link between the level of *RTNLB16* expression in the mutant and the severity of the growth phenotype.

To investigate whether enhanced *RTNLB16* expression alone can cause growth inhibition, we generated transgenic Arabidopsis lines overexpressing the genomic RTNLB16 fragment harboring isoforms 1-6 controlled by the CaMV35S promoter or by the β-estradiol-induced cassette in pMDC7 vector (Curtis and Grossniklaus, 2003). Despite high expression levels in leaves, no notable phenotype was observed in any of the transgenic lines growing under CLL or 16:8 conditions (Fig. S8), supporting the conclusion that the distinct phenotype of rtnlb16-1 was caused by a disturbance in RTNLB16 expression involving a combination of isoform 7 knockout and overactivation of the other isoforms.

RNA-seq comparisons between *rtnlb16-1* and the wild type revealed 980 and 3871 differentially expressed genes (DEGs) under continuous low light (CLL) or 16:8 conditions, respectively, evenly induced and suppressed (Fig. 7A, B). The number of DEGs between the mutant and the wild type, which was 3.95 times higher under 16:8 compared with CLL conditions, corresponded to a 5-fold difference in RTNLB16 transcript levels between the two conditions (Fig. 7A), suggesting a causal link. A Venn diagram comparison identified 351 CLL-exclusive DEGs, 629 DEGs common to CLL and 16:8, and 3242 16:8-exclusive DEGs (Fig. 7B). Gene Ontology (GO) enrichment analysis for biological processes in each DEGs cluster revealed distinct and common processes. The CLL-exclusive DEGs were enriched in transcripts for defense responses to biotic stress, response to salicylic acid (SA), and response to abscisic acid (ABA) (Fig. 7C, Table S2). Within the cluster common to CLL & 16:8, response to SA, defense response, and responses to biotic stress agents were much more pronounced than in the CLL-exclusive (Fig. 7D, Table S3). Other enriched processes included leaf senescence, photosynthesis, response to water deprivation, and responses to ABA and jasmonic acid (JA). The 16:8-exclusive DEGs cluster included many processes, among which the most pronounced were response to auxin, response to light, photosynthesis, response to wounding, response to JA, cell division, and tissue development (Fig. 7E, Table S4). Also, the response to ABA was enriched in this cluster. These results suggest that RTNLB16 is involved directly or indirectly in various activities, including responses to biotic and abiotic stress, as well as photosynthesis and development. Furthermore, enrichment in defense response and responses to SA and ABA, common to the three DEG clusters, suggests a direct involvement of RTNLB16 in these processes. Within the 16:8-exclusive DEGs between the mutant and the wild type, the expression of 509 genes was also different in the mutant between the two conditions, with near wild-type-like levels under CLL (Fig. 8A, B). It is, therefore, reasonable to infer that the transcriptional alteration within this cluster of 509 genes significantly contributed to the exacerbation of the mutant’s phenotype under 16:8 conditions. Significantly enriched processes within the 509 genes included defense responses, response to wounding, response to SA, and iron homeostasis (Fig. 8C, Table S5). Other processes included responses to auxin, JA, ABA, and leaf senescence.

**Fig. 7.**
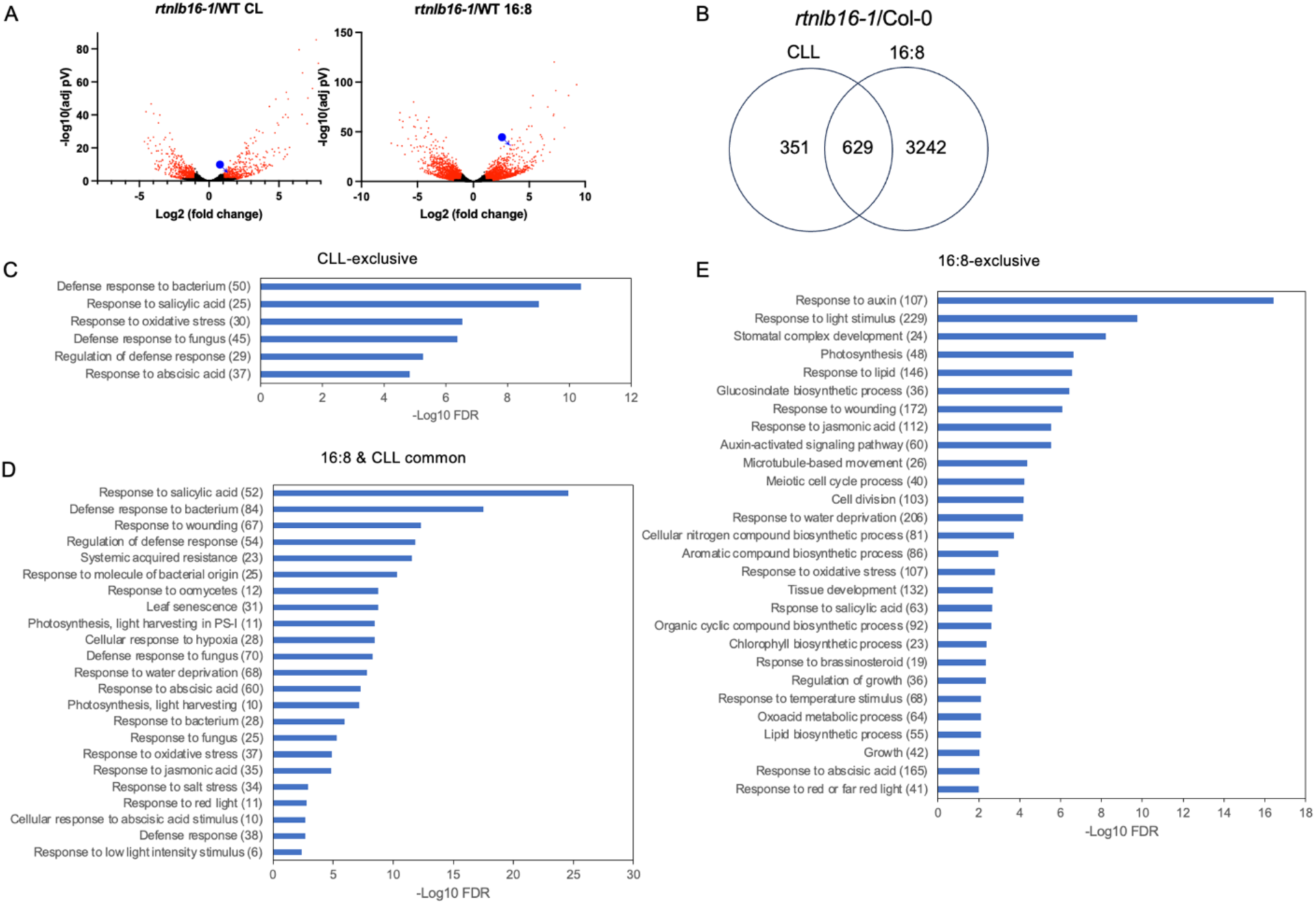
RNA seq comparison between rtnlb16-1 and the WT at CLL and 16:8 conditions. A. Volcano plots showing the dynamic range of DEGs distribution. The blue circles with the arrows point to the RTNLB16 DEG red dot value. B. Venn diagram of DEGs at CLL and 16:8 conditions, C-E, GO enrichment analysis for biological processes in each of the distinct sections in the Venn diagram.

**Fig. 8.**
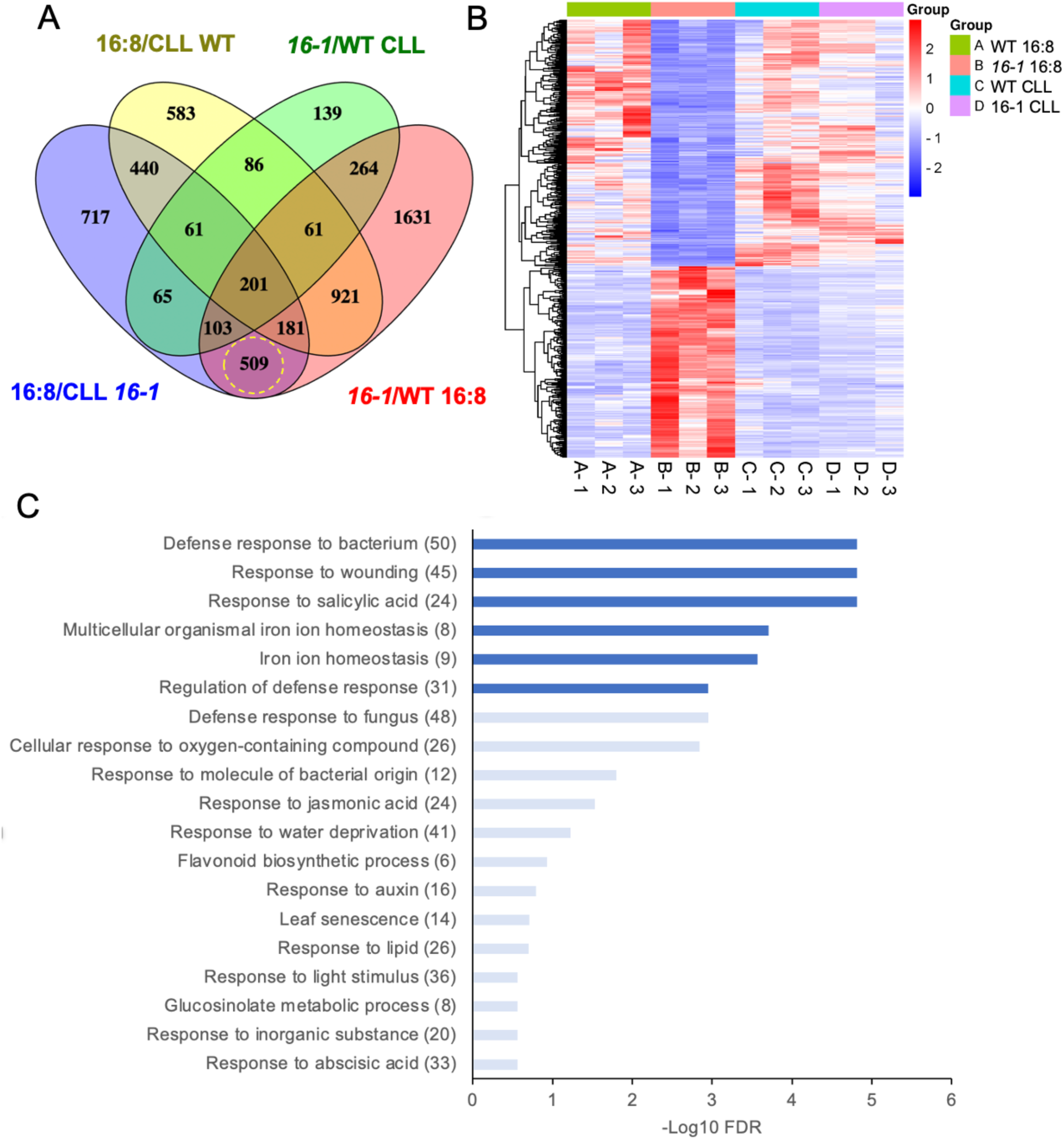
Four-way DEGs comparisons. A. Venn diagram of DEGs comparisons. The yellow dashed circle highlights the 509 DEGs in the mutant between conditions and at 16:8 between the mutant and WT. B. Heat map of the normalized expression values (RPKM) of the 509 DEGs (A) in all 12 RNAseq samples. C. GO enrichment analysis for biological processes in the 509 DEGs. Blue bars indicate GO terms with a false discovery rate (FDR; Benjamini) < 0.01. Light blue bars indicate Go terms FDR values below the threshold (i.e., FDR > 0.01) yet having pValue <0.01.

We then tested whether the responses to senescence stimulus or ABA were altered in the rtnlb16 mutants. Seedlings of WT, *rtnlb16-1*, and *rtnlb16-2* grown at CLL for 18 days were transferred to complete dark stress for seven days to induce senescence, after which they were returned to CLL for recovery. Results showed that while the WT and *rtnlb16-2* lost most of their chlorophyll during the extended dark period and were completely dead within 3 days after returning to CLL, *rtnlb16-1* survived and made a complete recovery (Fig. S9). This result suggested that the unique alteration in *RTNLB16* expression in this mutant led to senescence tolerance.

The response to ABA was evaluated using a seed germination assay, in which *rtnlb16* mutants were also compared with *abi4-1* mutant on media containing 0.3µM ABA. Results indicated increased insensitivity to ABA in both *rtnlb16* mutants, with around 60% germination compared to 20% and 90% germination for the wild type and *abi4-1*, respectively (Fig. 9). These results correspond with the transcriptional changes in ABA-responsive genes in *rtnl16-1* and support the involvement of RTNLB16 activity in ABA regulation.

**Fig. 9.**
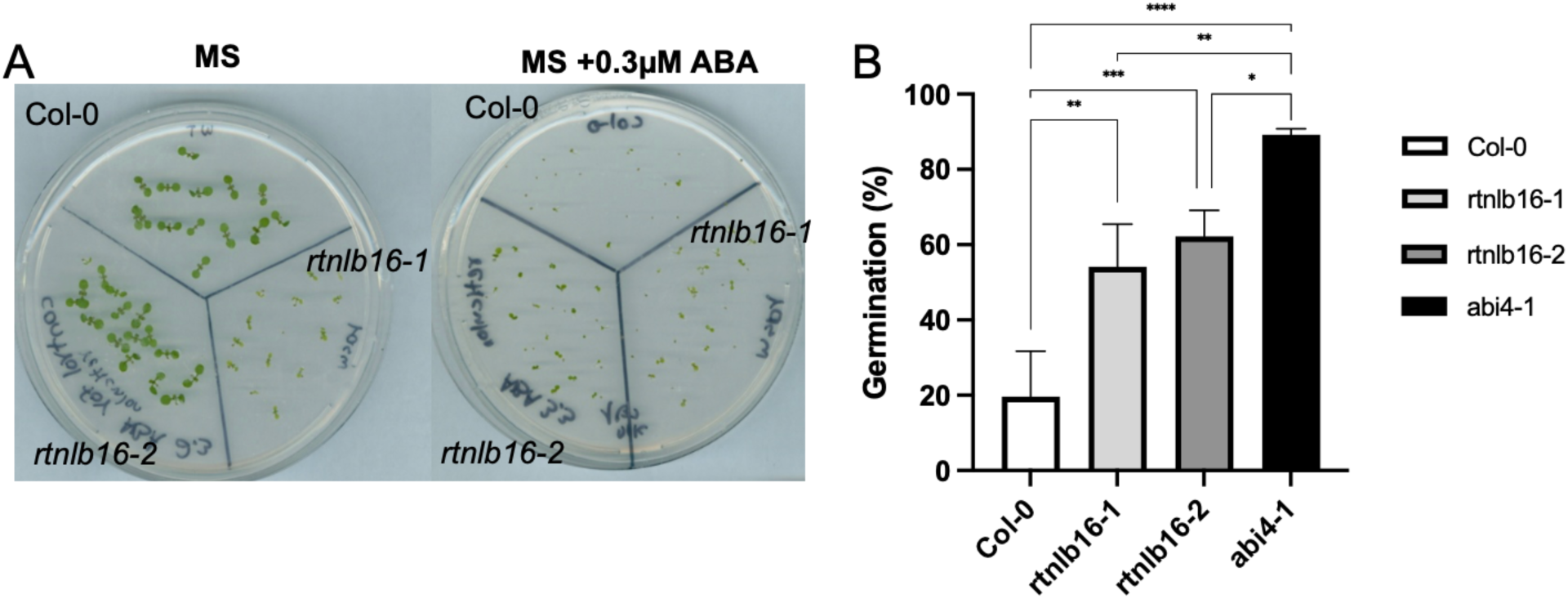
Increased insensitivity to ABA in rtnlb16 mutants. A. Representing photo showing the reduced sensitivity to germination of ABA in the mutants seven days after imbibition. B. Graph of germination score in the presence of 0.3 µM ABA 5 days after imbibition. *, **, ***, and **** indicate P<0.05, P<0.01, P<0.001, and P<0.0001, respectively in one-way ANOVA and Tukey’s multiple comparisons test.

## Discussion

In this study, we clarified that the T-DNA insertion in the Salk_122275 mutant (*rtnlb16-1*, formerly *oiwa-1*) is located within the RTNLB16 gene and not MSD1, as was erroneously indicated (Martin et al., 2013). Rectifying this mistake provided an opportunity to gain insight into the function of perhaps the most complex reticulon-like protein family member, having seven splice isoforms potentially coding for distinct protein variants. Our results suggest that all seven *RTNLB16* isoform transcripts are expressed in the wild type, whereas the rtnlb16-1 mutant is knocked out in the upstream ORF of isoform 7, while the expression of the other isoforms is overexpressed (Fig. 1).

Although the potential for gain-of-function in Salk lines is acknowledged (Ulker et al., 2008), very few studies have reported it. One such example is Salk_044777, in which the T-DNA insertion within the first exon of the NAC transcription factor AtNTL7 led to increased expression of a truncated form of the gene, contributing to ER stress resistance (Chi et al., 2017). Here, in the case of *rtnlb16-1*, a concatenated double-unversed T-DNA insert caused a distinct combination of loss- and gain-of-function within the same gene that manifested in dramatic inhibition of growth and chlorophyll accumulation. We suggest that this novel disturbance in *RTNLB16* expression altered the balance in the stoichiometry among the different splice forms of the gene in rtnlb16-1, impairing normal cellular activity. While high 35S promoter activity mediated by photoperiodic induction (16:8 conditions) increases disproportion among *RTNLB16* isoforms and leads to growth inhibition in the mutant, decreasing 35S promoter activity by continuous light (CLL conditions) lowers this imbalance and alleviates the phenotypic symptoms. Supporting this hypothesis is the finding that mere overexpression of *RTNLB16* or decreased expression with disruption to isoforms 1 and 3, as exists in *rtnlb16-2*, did not lead to growth inhibition.

Confocal microscopy with RTNLB16.5:GFP showed partial co-localization with the ER mCherry marker in the structures of the tubular ER network and additional intracellular localizations that may belong to the Golgi apparatus and PD (Fig. 4). These findings suggest that RTNLB16 may also be assigned to the inter- and intracellular trafficking pathways. In addition to shaping the ER membranes, mammalian reticulons (RTNs) function in trafficking material from the ER to the Golgi apparatus and the plasma membrane (Yang and Strittmatter, 2007; Di Sano et al., 2012). Still, the localization of RTNLB16 requires further investigation. PD formation involves extensive remodeling of the cortical ER into tightly furled desmotubules, axial structures that run through the PD pore connecting the ER of two adjacent cells (Ehlers and Kollmann, 2001). RTNLBs were suggested to function in the process that shapes and constricts the ER membranes into extremely fine desmotubules (Sparkes et al., 2010). RTNLB3 and 6 have been shown to reside in PD, in addition to the ER, and interact with many ER and PD proteins as well as several other RTNLB family members (Kriechbaumer et al., 2015; Knox et al., 2015; Fernandez-Calvino et al., 2011). Thus, these findings support our conclusion that RTNLB16 is also localized to PD.

The pleiotropic phenotype of the *rtnlb16-1* mutant under 16:8 conditions included short inflorescences and reduced seed production (Fig. S4). The heterozygous *rtnlb16*/RTNLB16 produced significantly fewer seeds than the homozygous mutant (Fig. 3), suggesting non-Mendelian inheritance of this allele. T-DNA mutants with increased seed abortion in the heterozygote are associated with T-DNA-mediated chromosomal translocation (Min et al., 2020; Curtis et al., 2009; Gang et al., 2019). Min et al. (2020) reported gametophytic abortion with distorted Mendelian segregation in heterozygote but not homozygote T-DNA mutant due to genome rearrangement, not because silencing of the impacted Anti-Silencing Function 1B (ASF1B) gene locus on Arabidopsis chromosome 5. Inversion of chromosomal fragments due to inter-chromosomal rearrangement leads to the suppression of recombination during homologous chromosomal pairing in Meiosis, resulting in resulted in gamete aneuploidy (Curtis et al., 2009). Multiple T-DNA insertions and concatenation of copies are common in plant transformation experiments, often leading to complex mutations, including flanking filler sequences and large-scale genomic rearrangements (Thomson et al., 2023). Translocation frequency surveyed in a collection of 64 independent Arabidopsis Salk lines detected chromosomal translocations in nearly 20% of the mutants, which in most cases led to abnormal pollen phenotype (Clark and Krysan, 2010). However, the fact that the concatenated T-DNA insertion at the RTNLB16 locus led to the deletion disrupting both RTNLB16 and MSD1 promoters, together with transcriptional activation at both sides of the insertion site (Figs.1, 6), suggests no translocation occurred in this locus (Fig. 1a). Yet, we cannot rule out genome rearrangement events in *rtnlb16-1* in other loci due to partial T-DNA insertions or unsuccessful integration events.

If no chromosomal translocation occurred in *rtnlb16-1*, an alternative explanation for the increased abortion in the heterozygotes may be directly linked to the disturbance in RTNLB16 expression and activity. The function of the ER is critical for nuclear fusion between the two polar nuclei during female gametogenesis as well as during double fertilization in the fusion between the sperm cells with the egg or central cell (Maruyama et al., 2014, 2020; Kobayashi and Nishikawa, 2023). Therefore, if RTNLB16 is involved in cellular activities in gametes, then asymmetry distribution of RTNLB16 isoforms between mutant pollen and WT ovules (and vice versa) might impair double fertilization. Additional experiments focusing on the reproduction of *rtnlb16* mutants are required to determine which of the above scenarios occurred.

The number of DEGs obtained in the RNAseq experiment was in correlation with the degree of the disturbance in *RTNLB16* expression and the severity of the phenotype, the majority of which (80%) were obtained under 16:8 conditions. Within the DEGs common to CLL and 16:8 conditions, the most pronounced GO-enriched processes were related to biotic stress responses, including bacteria, fungi, wounding, SA, and JA. The constitutive transcriptional changes in these pathways in *rtnlb16-1* point to the involvement of RTNLB16 in the secretory network employed in defense responses. The cortical ER network of plants is known to play multiple roles in protein trafficking (Bellucci et al., 2017; Robinson et al., 2015; Brandizzi, 2018) and response to pathogens (Pattison and Amtmann, 2009; Beck et al., 2012). Furthermore, several AtRTNLBs were shown to play a role in defense response, agrobacterium infection and interact with viral or immune response proteins (Kriechbaumer et al., 2015; Lee et al., 2011; Huang et al., 2018, 2021; Huang and Hwang, 2020).

In addition to changes in defense responses, the transcriptional profile of *rtnlb16-1* included constitutive alterations (i.e., common to CLL and 16:8) in gene expression associated with leaf senescence, salt stress, drought, and ABA responses. We further demonstrated that the alteration in *RTNLB16* expression in the *rtnlb16-1* mutant increased its tolerance to senescence-inducing treatment and decreased sensitivity to ABA during seed germination. Enhanced insensitivity of ABA in rtnlb16-2 further supports the role of RTNLB16 in ABA signaling. Thus, the involvement of RTNLB16 protein(s) in senescence and osmotic stress responses is likely mediated by ABA. Leaf senescence involves phytohormones-associated pathways that include ABA, SA, and JA. These phytohormones were suggested as cargo for autophagy-related ER to vacuole membrane container delivery and catabolism (Kulich and Žárský, 2014).

The activity of the ER may directly impact cellular ABA homeostasis by the action of BG1, the ER-localized β-glucosidase that releases active ABA from the inactive ABA glucosyl ester (ABA-GE) storages (Burla et al., 2013; Han et al., 2020; Ondzighi-Assoume et al., 2016). Also, the SA level in the cell and the ER-stress response are closely associated, which may reciprocally impact each other (Wang et al., 2005; Liu et al., 2023; Poór et al., 2019). Therefore, balanced expression of the different RTNLB16 isoforms may be critical for proper ABA-, SA-, and JA-activated processes.

The alteration in the expression of the 509 genes observed in the *rtnlb16-1* mutant under photoperiodic conditions (Fig. 8) may be a consequence of the increased imbalance within *RTNLB16* expression. Misexpression of these genes, enriched for defense responses, SA, JA, auxin, and ABA, might have contributed the most to the exacerbation of the growth inhibition in the mutant under 16:8 conditions; activation of numerous genes associated with both biotic and abiotic stress responses in *rtnlb16-1* may have increased the energetic demand for defense and acclimation processes, diverting resources away from growth. The ER functions as a central hub for integrating signals from biotic and abiotic stress responses (Park and Park, 2019). RTNLB16 might play a crucial role in the ER network, coordinating the crosstalk between defense and acclimation responses when the plant faces a combination of biotic and abiotic stresses.

This study presents an Arabidopsis mutant line with a T-DNA insertion causing a unique aberration in *RTNLNB16* expression, including a knockout of an upstream ORF of one splice form while increasing the expression of the others. This alteration can be tuned by changing the diurnal light regime. More than 700,000 T-DNA lines were generated in different T-DNA insertion mutant projects in dicots and monocots (Jupe et al., 2019). With the majority of these T-DNA lines used for identifying gene silencing mutations (Ulker et al., 2008), events of gene activation or a combination of both, as in the case of *rtnlb16-1*, are often overlooked. Here, we demonstrate that by shifting plants between photoperiodic or continuous light conditions, one can easily distinguish between visible phenotypes resulting from transcriptional silencing, over-expression, or a combination of both in T-DNA lines harboring the 35S promoter. This simple conditional treatment test can be readily applied to uncover hidden gene functions in T-DNA line projects such as Salk and Gabi_kat, and revisit abandoned/discontinued mutants whose phenotypes could not be explained.

## Materials and Methods

### Plant Materials, Growth Conditions, and Treatments

All lines utilized in this study were of Arabidopsis thaliana, Columbia ecotype. RTNLB16 T-DNA insertion lines (*rtnlB16-1* - Salk_122275; *rtnlb16-2* – Salk_020022) were obtained from the Arabidopsis Biological Resource Center (https://abrc.osu.edu/). Homozygous lines were PCR-identified following SIGnAL Laboratory recommendations (http://signal.salk.edu/tdnaprimers.2.html).

Plants were cultivated in growth chambers (Percival AR-66; Percival Scientific) or temperature-controlled growth rooms at 23°C under a long day 16-/8-h light/dark cycle (80-100 µmol m^−2^ s^−1^), or continuous low light (CLL; 40 µmol m^−2^ s^−1^) photosynthetic flux density, and with 70% relative humidity. Due to rtnlb16-1’s extreme sensitivity to photoperiodic conditions, germination and initial growth occurred under CLL conditions in most experiments involving exposure to long-day conditions, unless noted otherwise.

### ABA Germination Assay

Freshly harvested seeds were surface sterilized and sown on 0.5 MS agar with or without 0.3µM ABA. Germination (radicle emergence) was assessed 5 days after imbibition under CLL conditions.

### Dark Stress

Two-week-old plants on MS agar plates growing under CLL were covered with aluminum foil for complete darkness and kept in a growth chamber for 7 days.

### Molecular Analyses

RNA extraction from fresh leaves was conducted with the TRIZOL reagent (Life Technologies) following the manufacturer’s recommendations. PCRs, cDNA synthesis, quantitative real-time PCR (qRT-PCR), and semi-quantitative PCRs were performed as previously described (Chen et al., 2014, 2021). The primer names and sequences are listed in Supplemental Table S1.

### SODs Activity Gel and Immunoblot

Leaf extracts were prepared from 250 mg leaves harvested from 3-week-old CLL-grown plants. Protein samples of 30 µg were separated on a 10% non-denaturing polyacrylamide gel, and SOD activity assays were performed according to Chu et al. 2005. Western blot was conducted as described by Chen 2021 with polyclonal anti-MSD1 antibodies (Agrissera, AS09 524) at a 1:3000 dilution.

### Cloning and Generation of Transgenic Plants

To determine the subcellular localization of RTNLB16, spliced variant (isoform) 5 CDS was PCR-amplified and inserted into Gateway^TM^ pDONOR221 entry vector and subsequently into the destination pH7FWG2 binary vector (Karimi et al., 2002), where it was fused to C-terminal GFP under the CaMV 35S promoter. Similarly, RTNLB16 isoform 5, as well as the genomic fragment of RTNLB16 harboring isoforms 1-6 (ATG to TGA), were cloned into pH7FWG2 and estrogen-inducible pMDC7 (Curtis and Grossniklaus, 2003) destination vectors. PCR amplification was carried out using Herculase II Fusion DNA Polymerase enzyme (Agilent Genomics, USA) with specific primers (Supplemental Table S1).

### Transient Expression in *N. benthamiana*

Agrobacterium infiltration was performed according to (Sparkes et al., 2006). In the co-expression experiments, the RTNLB16:GFP and ER-rk agrobacteria were mixed in a 1:1 ratio.

### Delayed Fluorescence

Chlorophyll (Chl) was evaluated in the entire rosette of 14-day-old plants by measuring delayed fluorescence using the NightSHADE LB985 imager (Berthold Technologies, Bad Wildbad, Germany) according to Gould et al. (2009)-19638147.

### Microscopy and Imaging

Fluorescence imaging in Agrobacterium-infiltrated *N. Benthamiana* leaf sections and transgenic Arabidopsis leaves was conducted using the Olympus-FV1000 confocal microscope, or the CRI Maestro II (PerkinElmer, USA) live imaging system for GFP imaging in 35S:GFP Arabidopsis seedlings.

### RNAseq and Bioinformatics

The library preparation procedure followed the protocol previously described by (Rutley et al., 2021). Twelve samples were sequenced on a single lane of an Illumina HiSeq 2500 sequencer in high-output mode, producing 30 to 50 million reads per sample. The data were aligned to the annotated TAIR10 (araport-11 release 06_2016) using STAR version 2.5.0a (Dobin et al., 2013). HTSeq (version 0.6.1p1) was employed to generate the count matrix (Anders et al., 2015). Differential expression analysis was conducted using DESeq2 (Love et al., 2014). Comparisons were made as follows:

Three replicates from sample A (Col-0 16:8) with the three replicates from sample B (*rtnlB16-1* 16:8).

Three replicates from sample C (Col-0 CLL) with the three replicates from sample D (*rtnlb16-1* CLL).

Three replicates from sample type A with the three replicates from sample type B. Three replicates from sample type C with the three replicates from sample type D.

Differentially expressed genes (DEGs) were defined as those exceeding a one log-fold change (p-value<0.01). The RNAseq data have been deposited at https://www.ncbi.nlm.nih.gov/geo/ with accession number GSE242257. Functional gene analysis was conducted using DAVID Bioinformatics Resources 6.8 (Sherman et al., 2022). Visualization of the results was performed with the Integrative Genomics Viewer (IGV) (Thorvaldsdóttir et al., 2013).

### Statistics

Graphs and statistical analyses for significant differences between mean values using one/two-way ANOVA and Tukey’s were conducted using GraphPad Prism v.10 software.

## Acknowledgments

We thank Yuval Nagar and Rotem Rosh who assisted in the research work during their undergraduate lab project.

## Supplementary data

Fig. S1. Salk_122275/*oiwa-1*/*rtnlb16-1* is not deficient in MSD1.

Fig. S2. Genotyping of *rtnlb16* T-TDNA mutants.

Fig. S3. Genomic sequence viewer of AT3G10915 and AT3G10920 indicating the correct and exact localization of the T-DNA inserts in the lines indicated

Fig. S4. Heterozygous Salk_122275 (*rtnlb16-1*) is indistinguishable from the wild type at the vegetative stages.

Fig. 5S. Co-localization of RTNLB16.5:GFP and the ER-mCherry marker in N. Benthamiana epidermis.

Fig. S6. Impact of short-term transition from CLL to 16:8 on *rtnlb16-1* plants.

Fig. S7. Light-dependent growth phenotype of mature flowering *rtnlb16-1*.

Fig. S8. Overexpression of RTNLB16 does not impact growth.

Fig. S9. Dark stress resistance in *rtnlb16-1* plants. Table S1. Primer’s list

Table S2. GO biological Processes enrichment for CLL-exclusive DEGs

Table S3. GO biological Processes enrichment for 16:8 and CLL-common DEGs.

Table S4. GO biological Processes enrichment for 16:8-exclusive DEGs.

Table S5. GO Biological Processes enrichment for 509 DEGs in Fig. 8.

## Author Contributions

T.K. and D.L. conducted experiments and analyses. H.B.A conducted bioinformatic data analysis of the RNA seq, and G.M. Conceived the study, designed the experiments, wrote the manuscript, and prepared figures.

