## Supplemental data for "Disproportion among reticulon-like 16 (RTNLB16) splice variants disrupts growth and decreases sensitivity to ABA and senescence in Arabidopsis"

### Supplementary Figures

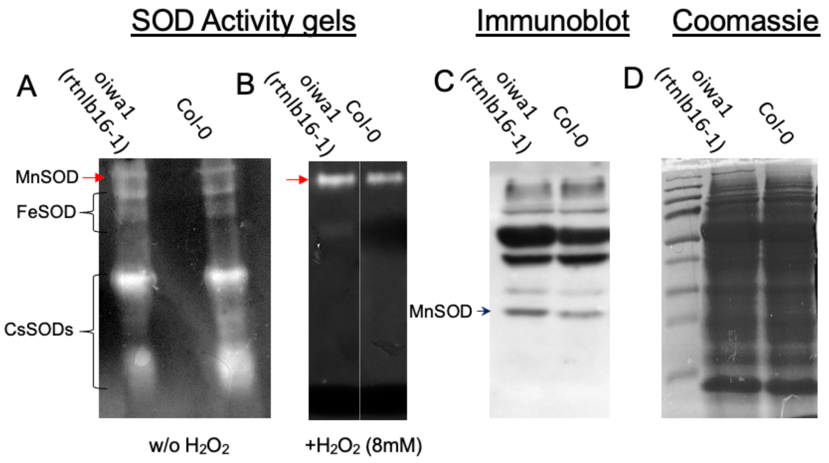


**Fig. S1. Salk_122275/*oiwa-1*/*rtnlb16-1* is not deficient in MSD1.** A and B, SOD activity gels, B, plus H2O2 treatment that inactivates all other SODs but MSD1. C, western blot. D. Coomassie loading control.

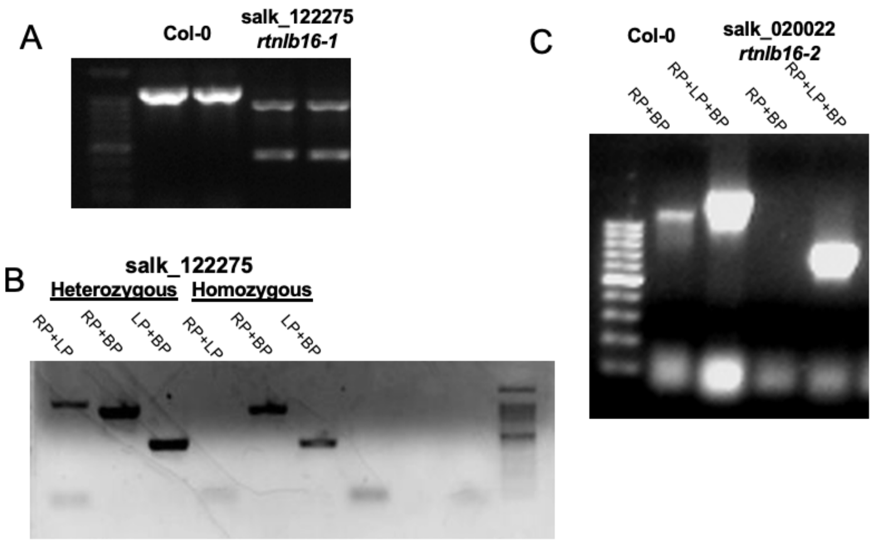


**Fig. S2. Genotyping of *rtnlb16* T-TDNA mutants.** A) PCR products using three primers: Salk_122275 LP+RP + LBb1 (left border primer) showing two products on homozygous Salk_122275 gDNA that are lower than the WT ban, indicating a double inverted repeat insertion. B. Two primers PCR Genotyping of Salk_122275 homozygous and heterozygous plant. C. PCR products using two or three primers:

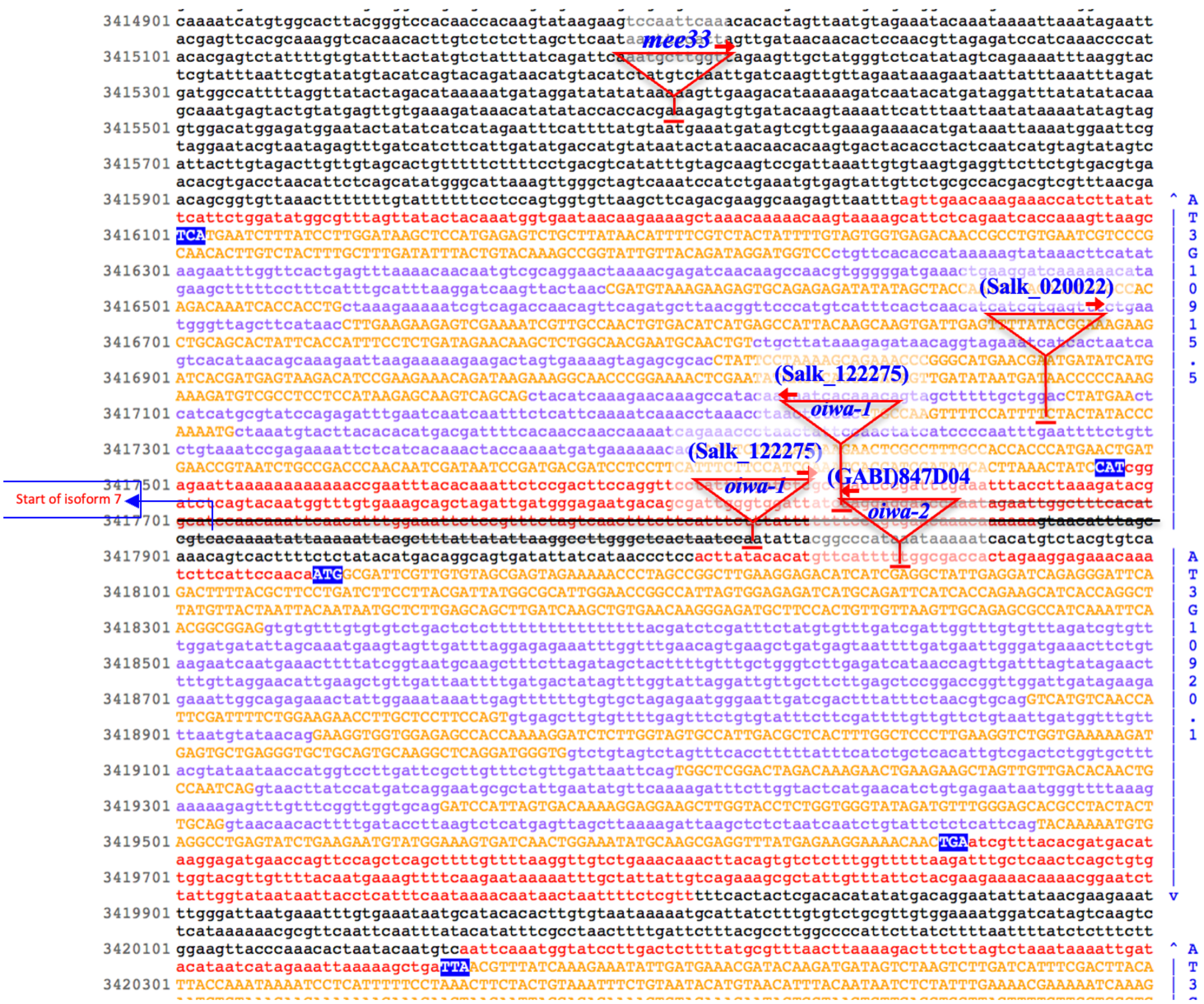


**Fig. S3. Genomic sequence viewer of AT3G10915 and AT3G10920 indicating the correct and exact localization of the T-DNA inserts in the lines indicated in this paper and in Martin et al. 2013**. The black strikethrough line indicated the genomic segment deleted in Salk_122275 from the integration event of the T-DNAs. The sequence of AT3G10915 is presented according to slice variant (isoform) 5. Exons = Upper case orange, introns = purple, UTRs = red, intergenic/promoters/UTR (of other isoforms) = black letters. The transcriptional start site of isoform 7 is indicated. The location of oiwa-2 and mee33, mentioned by Martin et al. 2013, is also indicated.

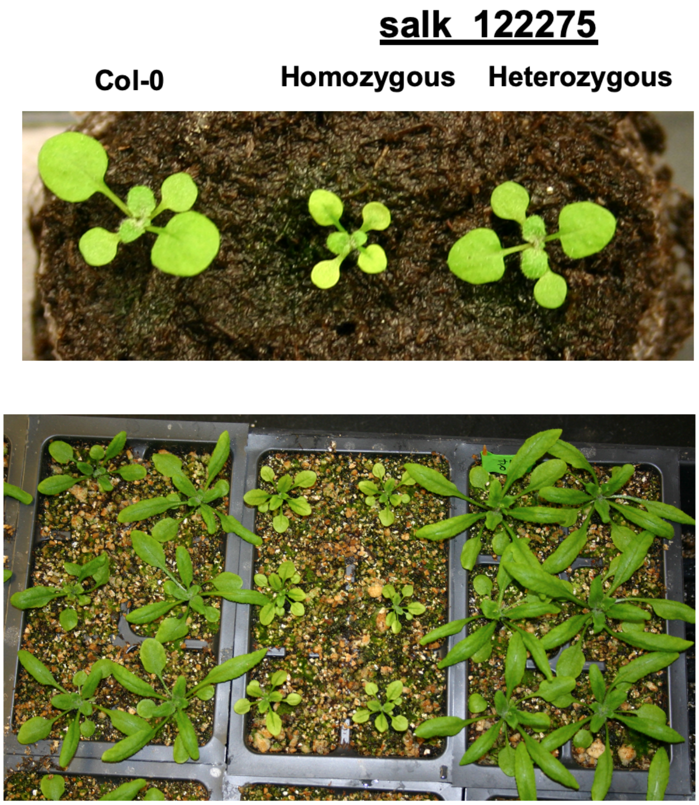


**Fig. S4. Heterozygous Salk_122275 (*rtnlb16-1*) is indistinguishable from the wild type at the vegetative stages.** Plants were grown under CLL conditions. Top, two-week-old plants. Bottom, four-weeks-old plants.

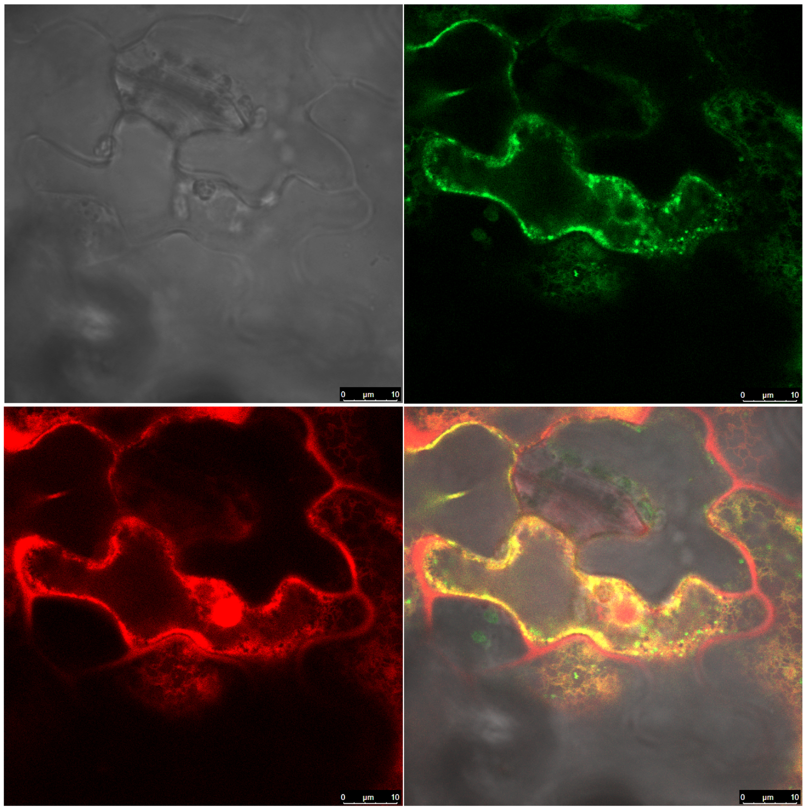

**Fig. 5S. Co-localization of RTNLB16.5:GFP and the ER-mCherry marker in N. Benthamiana epidermis.**

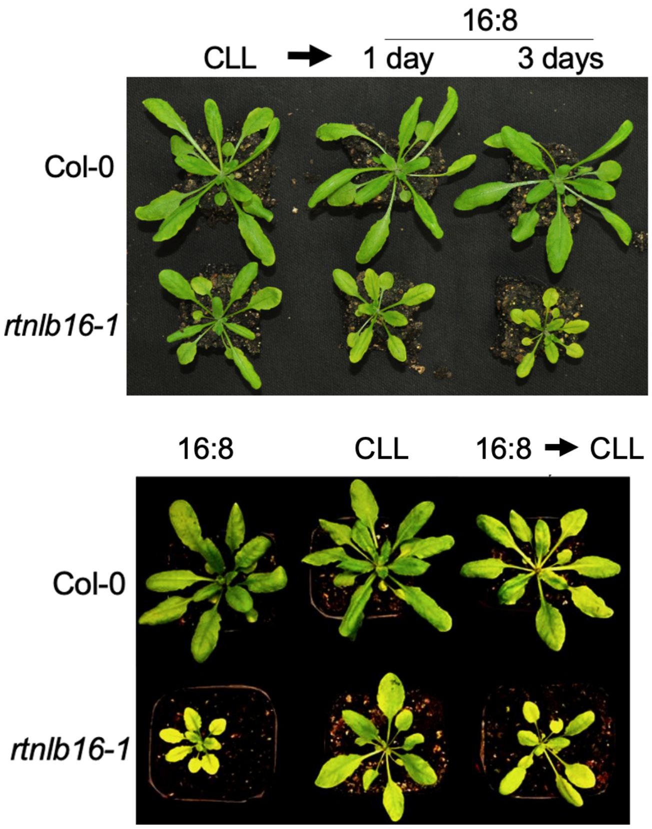


**Fig. S6. Impact of short-term transition from CLL to 16:8 on *rtnlb16-1* plants.** Three weeks old plants after 1 or 3 days under 16:8 conditions. Plants from this experiment were used for RNA extraction and semi-qRT PCR presented in Fig. 6B.

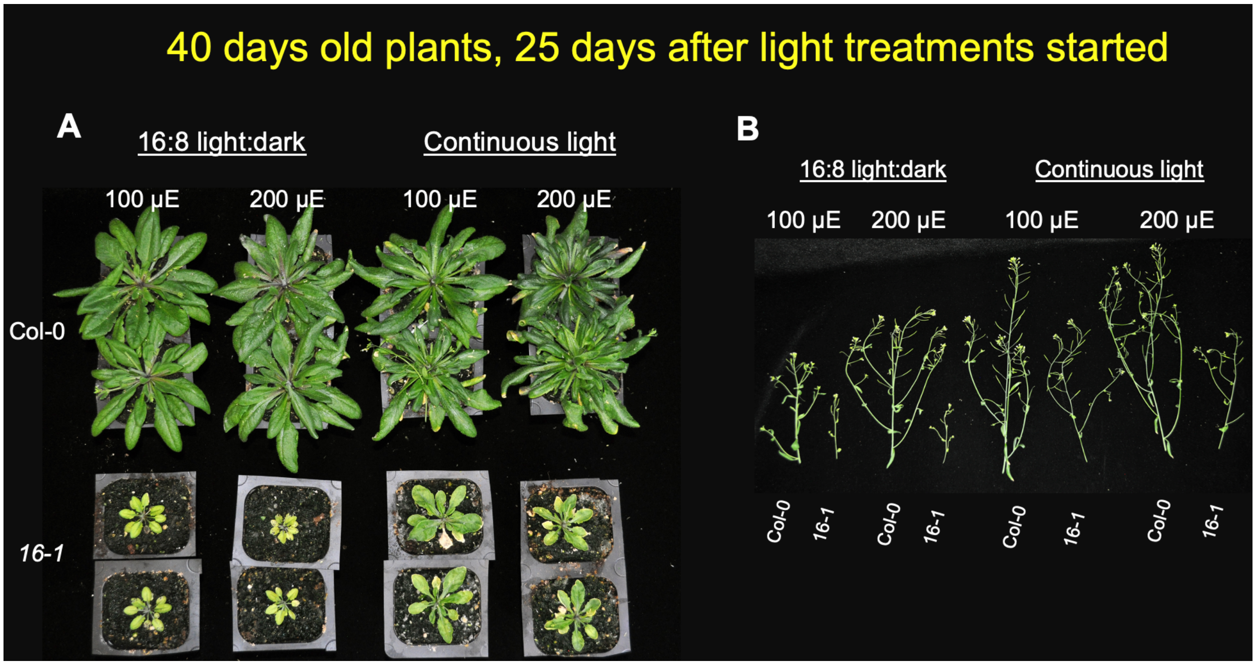


**Fig. S7. Light-dependent growth phenotype of mature flowering *rtnlb16-1*.** A. Two weeks old plants grown under CLL were transferred to continuous light (CL) or 16:8 conditions with a moderate light intensity of either 100 or 200 µE (µmol*$S^{-1}$*$m^{-2}$). A. Photos of the plants described in Fig. 5B, C 10 days later under the corresponding light treatments. The inflorescence stems presented in B, were cut for clear capturing of the rosettes.

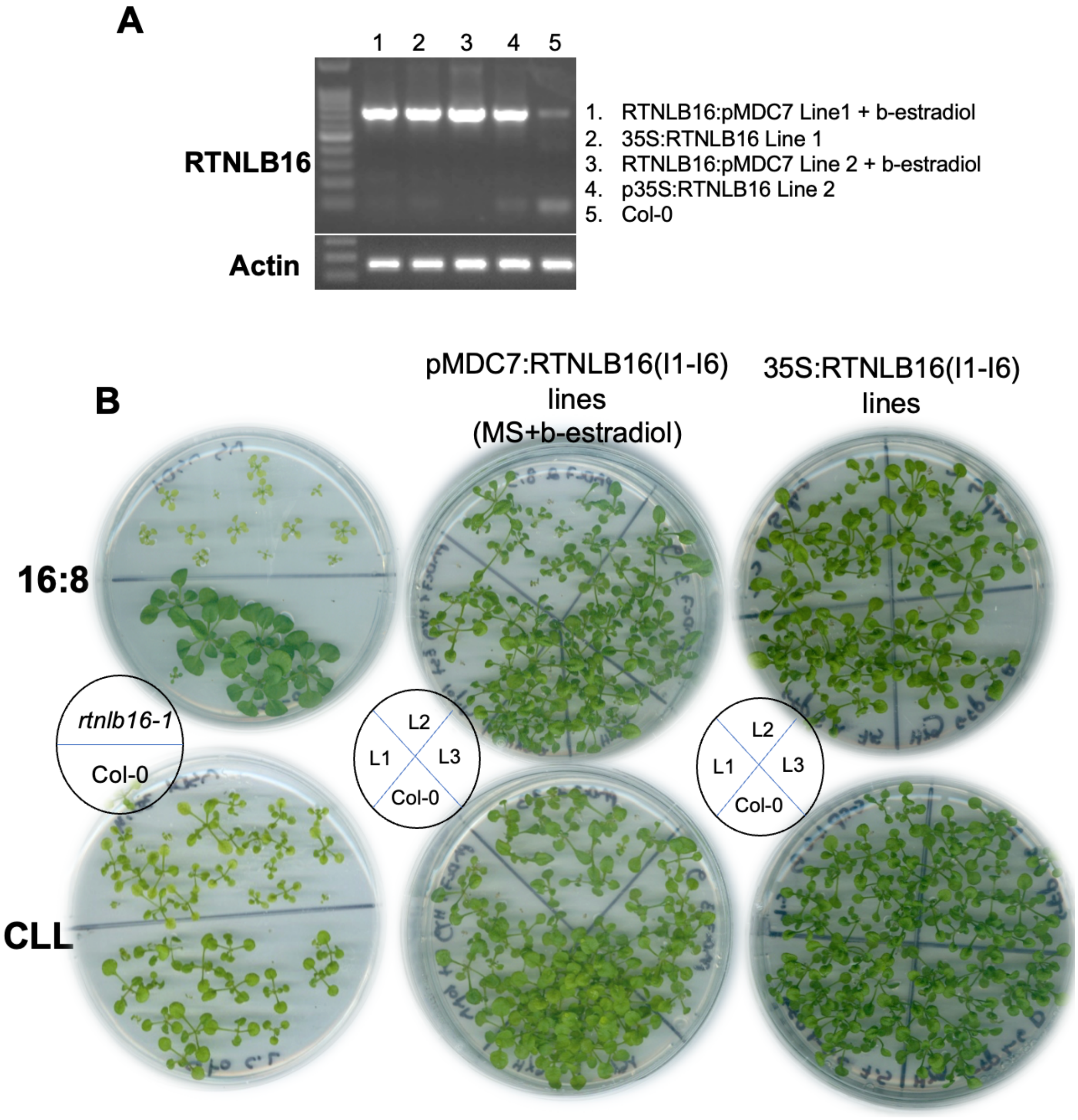


**Fig. S8. Overexpression of RTNLB16 does not impact growth.** Transgenic line of RTNLB16 gene harboring the reading frames of isoforms 1-6 controlled by estrdiol-induced cassette or CaMV 35S promoter. A. PCR amplification of RTNLB16. B. Growth of two-week-old transgenic plants grown under CLL or 16:8 conditions compared to rtnlb16-1 and WT. Three independent lines (L1-3) of each transgenic type were grown next to Col-0 in a plate.

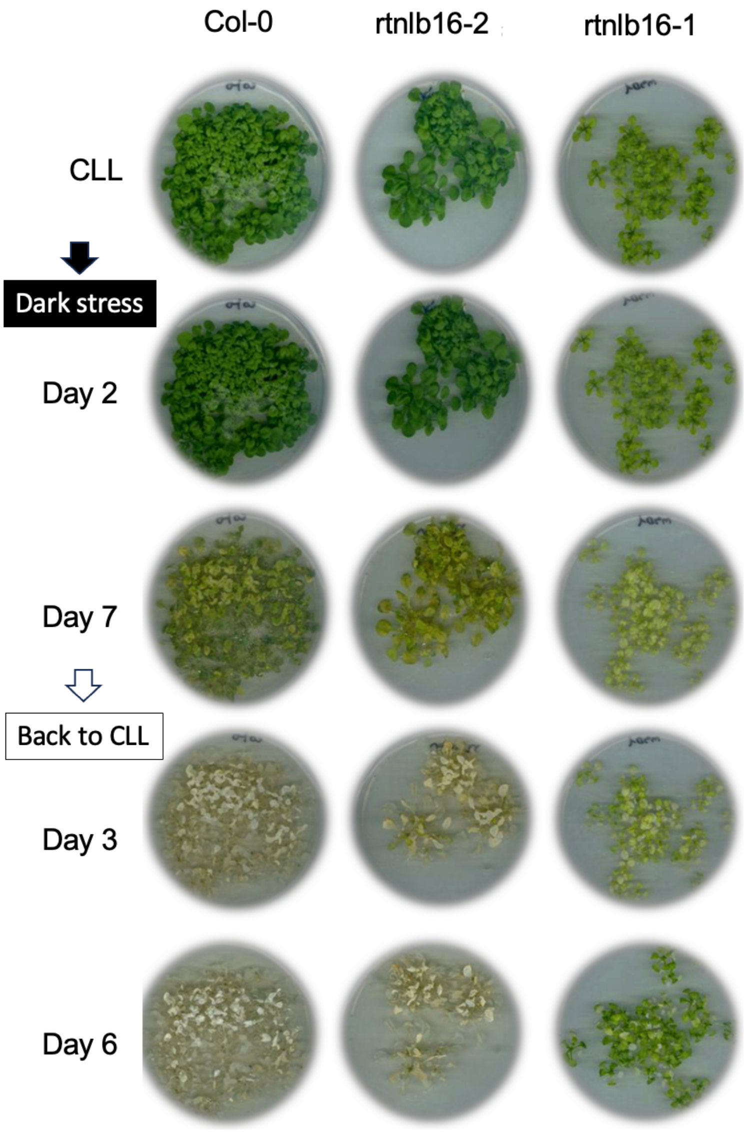


**Fig. S9. Dark stress resistance in *rtnlb16-1* plants**. Eighteen days old plants grown at CLL were transferred to complete darkness for seven days (temporarily taken out for capturing photos) followed by recovery back at CLL conditions.

#### Supplementary table

| Primer name | Target gene | Purpose | Sequence | Products&sizes | Relevant Fig. |
|---|---|---|---|---|---|
| P1 (For) | RTNLB16 | sq-PCR | 5' ATGCATGTGAAAGCCAATTC 3' | | |
| P2 (Rev) | RTNLB16 | sq-PCR | 5' CACCACCCATGAACTGATGA 3' | I1-5=320 bp, I7=170bp | Fig. 1E |
| P3 (For) | RTNLB16 | sq-PCR | 5' ATCGCTGTCATTCTCCCATC 3' | I1= 876 bp, I2= 773 bp, I3=1113 bp, I4=777bp, | |
| P4 (Rev) | RTNLB16 | sq-PCR | 5' TGTAGTGGTGAGACAACCGC 3' | I5=842 bp, I6=677 bp, I7=641 bp, gDNA=1490 | Fig. 1F |
| P5 (For) | RTNLB16 | sq-PCR | 5' GCTAACCCATTCAGAAACTCACG 3' | | |
| P4 (Rev) | RTNLB16 | sq-PCR | 5' TGTAGTGGTGAGACAACCGC 3' | I1=266 bp, gDNA=443 bp | Fig.1F |
| RTNLB16qRT_For | RTNLB16 | qRT-PCR | 5' GGAGGAGGCGACATCTTTCT 3' | I1-7=142 | Figs.1G,6B |
| RTNLB16qRT_Rev | RTNLB16 | qRT-PCR | 5' CCGGGCATGAACGAATGATA 3' | | |
| MSD1_F | MSD | qRT-PCR | 5' CGATTCGTTGTGTAGCGAGTAG 3' | | Fig.1G |
| MSD1_R | MSD | qRT-PCR | 5' GATGCTTCTGGTGATGAATCTG 3' | | |
| ACT2_F | Actin-2 | | GTCGTACAACCGGTATTGTGCTG | | Fig.1G |
| ACT2_R | Actin-2 | | CCTCTCTCTGTAAGGATCTTCATGAG | | |
| Salk_122275 LP | At3g10915/at3g10920 | Genotyping | 5' CGATCCTCCTTCATTTCTTCC 3' | | |
| Salk_122275 RP | At3g10915/at3g10920 | Genotyping | 5' TGATCTCAAGACCCAGCAAAC 3' | | |
| SALK_020022 | At3g10915 | | 5' AAGCCACAGACAAATCACCAC 3' | | |
| SALK_020022 | At3g10915 | | 5' TTCCTCCTTGATAATCCACCC 3' | | |

**Table S1. Primer's list**

| Category | Term | Count | % | PValue | Benjamini | FDR |
|---|---|---|---|---|---|---|
| GOTERM_BP_DIRECT | GO:0042742~defense response to bacterium | 50 | 14.36782 | 7.06E-14 | 4.23E-11 | 4.17E-11 |
| GOTERM_BP_DIRECT | GO:0009751~response to salicylic acid | 25 | 7.183908 | 3.22E-12 | 9.66E-10 | 9.51E-10 |
| GOTERM_BP_DIRECT | GO:0006979~response to oxidative stress | 30 | 8.62069 | 1.46E-09 | 2.92E-07 | 2.88E-07 |
| GOTERM_BP_DIRECT | GO:0050832~defense response to fungus | 45 | 12.93103 | 2.83E-09 | 4.24E-07 | 4.18E-07 |
| GOTERM_BP_DIRECT | GO:0031347~regulation of defense response | 29 | 8.333333 | 4.62E-08 | 5.53E-06 | 5.45E-06 |
| GOTERM_BP_DIRECT | GO:0009737~response to abscisic acid | 37 | 10.63218 | 1.53E-07 | 1.53E-05 | 1.51E-05 |
| GOTERM_BP_DIRECT | GO:0014070~response to organic cyclic compound | 15 | 4.310345 | 2.39E-04 | 0.020452 | 0.020144 |
| GOTERM_BP_DIRECT | GO:0009753~response to jasmonic acid | 19 | 5.45977 | 3.70E-04 | 0.027685 | 0.027269 |
| GOTERM_BP_DIRECT | GO:0009617~response to bacterium | 14 | 4.022989 | 4.43E-04 | 0.029501 | 0.029058 |
| GOTERM_BP_DIRECT | **GO:0030968~endoplasmic reticulum unfolded protein response** | 5 | 1.436782 | 8.37E-04 | 0.045572 | 0.044888 |
| GOTERM_BP_DIRECT | GO:0031348~negative regulation of defense response | 5 | 1.436782 | 8.37E-04 | 0.045572 | 0.044888 |
| GOTERM_BP_DIRECT | GO:0009611~response to wounding | 25 | 7.183908 | 0.001253 | 0.06253 | 0.06159 |
| GOTERM_BP_DIRECT | GO:0009627~systemic acquired resistance | 7 | 2.011494 | 0.003636 | 0.167541 | 0.165024 |
| GOTERM_BP_DIRECT | GO:0002237~response to molecule of bacterial origin | 8 | 2.298851 | 0.004588 | 0.184265 | 0.181496 |
| GOTERM_BP_DIRECT | GO:0006970~response to osmotic stress | 14 | 4.022989 | 0.004614 | 0.184265 | 0.181496 |
| GOTERM_BP_DIRECT | GO:0042435~indole-containing compound biosynthetic process | 5 | 1.436782 | 0.007994 | 0.299287 | 0.29479 |
| GOTERM_BP_DIRECT | GO:0009408~response to heat | 11 | 3.16092 | 0.008631 | 0.30411 | 0.29954 |
| GOTERM_BP_DIRECT | GO:0006749~glutathione metabolic process | 5 | 1.436782 | 0.011776 | 0.37998 | 0.374271 |
| GOTERM_BP_DIRECT | GO:0009414~response to water deprivation | 27 | 7.758621 | 0.012053 | 0.37998 | 0.374271 |
| GOTERM_BP_DIRECT | GO:0009620~response to fungus | 10 | 2.873563 | 0.014306 | 0.428478 | 0.42204 |
| GOTERM_BP_DIRECT | GO:0010150~leaf senescence | 10 | 2.873563 | 0.015229 | 0.431515 | 0.425032 |
| GOTERM_BP_DIRECT | GO:0071310~cellular response to organic substance | 8 | 2.298851 | 0.015849 | 0.431515 | 0.425032 |
| GOTERM_BP_DIRECT | GO:0009625~response to insect | 4 | 1.149425 | 0.018472 | 0.481069 | 0.473841 |
| GOTERM_BP_DIRECT | GO:0090558~plant epidermis development | 10 | 2.873563 | 0.0229 | 0.562751 | 0.554296 |
| GOTERM_BP_DIRECT | GO:0009407~toxin catabolic process | 4 | 1.149425 | 0.023495 | 0.562751 | 0.554296 |
| GOTERM_BP_DIRECT | GO:0010035~response to inorganic substance | 14 | 4.022989 | 0.024427 | 0.562751 | 0.554296 |
| GOTERM_BP_DIRECT | GO:0010015~root morphogenesis | 11 | 3.16092 | 0.029539 | 0.63237 | 0.622868 |
| GOTERM_BP_DIRECT | GO:0098542~defense response to other organism | 16 | 4.597701 | 0.02956 | 0.63237 | 0.622868 |
| GOTERM_BP_DIRECT | GO:0006952~defense response | 18 | 5.172414 | 0.034333 | 0.675734 | 0.665581 |
| GOTERM_BP_DIRECT | GO:1901701~cellular response to oxygen-containing compound | 13 | 3.735632 | 0.034346 | 0.675734 | 0.665581 |
| GOTERM_BP_DIRECT | GO:0008643~carbohydrate transport | 5 | 1.436782 | 0.034971 | 0.675734 | 0.665581 |
| GOTERM_BP_DIRECT | GO:0051707~response to other organism | 3 | 0.862069 | 0.040483 | 0.757783 | 0.746397 |
| GOTERM_BP_DIRECT | GO:0080027~response to herbivore | 3 | 0.862069 | 0.043623 | 0.79182 | 0.779923 |
| GOTERM_BP_DIRECT | GO:0071456~cellular response to hypoxia | 8 | 2.298851 | 0.04553 | 0.801811 | 0.789764 |
| GOTERM_BP_DIRECT | GO:1901617~organic hydroxy compound biosynthetic process | 3 | 0.862069 | 0.04685 | 0.801811 | 0.789764 |
| GOTERM_BP_DIRECT | GO:0070212~protein poly-ADP-ribosylation | 2 | 0.574713 | 0.052387 | 0.865309 | 0.852308 |
| GOTERM_BP_DIRECT | GO:0006468~protein phosphorylation | 19 | 5.45977 | 0.05345 | 0.865309 | 0.852308 |
| GOTERM_BP_DIRECT | GO:2000280~regulation of root development | 4 | 1.149425 | 0.056309 | 0.887611 | 0.874274 |
| GOTERM_BP_DIRECT | GO:0006457~protein folding | 6 | 1.724138 | 0.062651 | 0.943898 | 0.929716 |
| GOTERM_BP_DIRECT | GO:0015692~lead ion transport | 2 | 0.574713 | 0.06505 | 0.943898 | 0.929716 |
| GOTERM_BP_DIRECT | GO:0009415~response to water | 5 | 1.436782 | 0.065151 | 0.943898 | 0.929716 |
| GOTERM_BP_DIRECT | GO:0016310~phosphorylation | 8 | 2.298851 | 0.066183 | 0.943898 | 0.929716 |
| GOTERM_BP_DIRECT | GO:0043412~macromolecule modification | 9 | 2.586207 | 0.071767 | 0.999734 | 0.984712 |
| GOTERM_BP_DIRECT | GO:0071695~anatomical structure maturation | 5 | 1.436782 | 0.085005 | 1 | 0.986622 |
| GOTERM_BP_DIRECT | GO:1900367~positive regulation of defense response to insect | 2 | 0.574713 | 0.089873 | 1 | 0.986622 |
| GOTERM_BP_DIRECT | GO:0072334~UDP-galactose transmembrane transport | 2 | 0.574713 | 0.089873 | 1 | 0.986622 |
| GOTERM_BP_DIRECT | GO:0009651~response to salt stress | 14 | 4.022989 | 0.095523 | 1 | 0.986622 |

**Table S2. GO biological Processes enrichment for CLL-exclusive DEGs**.

| Category | Term | Count | % | PValue | Benjamini | FDR |
|---|---|---|---|---|---|---|
| GOTERM_BP_DIRECT | GO:0009751~response to salicylic acid | 52 | 8.4006462 | 3.99E-28 | 2.93E-25 | 2.71E-25 |
| GOTERM_BP_DIRECT | GO:0042742~defense response to bacterium | 84 | 13.570275 | 9.55E-21 | 3.51E-18 | 3.25E-18 |
| GOTERM_BP_DIRECT | GO:0009611~response to wounding | 67 | 10.82391 | 2.04E-15 | 5.00E-13 | 4.62E-13 |
| GOTERM_BP_DIRECT | GO:0031347~regulation of defense response | 54 | 8.723748 | 8.36E-15 | 1.54E-12 | 1.42E-12 |
| GOTERM_BP_DIRECT | GO:0009627~systemic acquired resistance | 23 | 3.7156704 | 2.18E-14 | 3.21E-12 | 2.96E-12 |
| GOTERM_BP_DIRECT | GO:0002237~response to molecule of bacterial origin | 25 | 4.0387722 | 4.02E-13 | 4.93E-11 | 4.55E-11 |
| GOTERM_BP_DIRECT | GO:1901701~cellular response to oxygen-containing compound | 43 | 6.9466882 | 2.34E-12 | 2.46E-10 | 2.27E-10 |
| GOTERM_BP_DIRECT | GO:0010150~leaf senescence | 31 | 5.0080775 | 2.12E-11 | 1.94E-09 | 1.79E-09 |
| GOTERM_BP_DIRECT | GO:0002239~response to oomycetes | 12 | 1.9386107 | 2.37E-11 | 1.94E-09 | 1.79E-09 |
| GOTERM_BP_DIRECT | GO:0010035~response to inorganic substance | 42 | 6.7851373 | 5.48E-11 | 3.55E-09 | 3.28E-09 |
| GOTERM_BP_DIRECT | GO:0071456~cellular response to hypoxia | 28 | 4.5234249 | 5.76E-11 | 3.55E-09 | 3.28E-09 |
| GOTERM_BP_DIRECT | GO:0009768~photosynthesis, light harvesting in photosystem I | 11 | 1.7770598 | 5.79E-11 | 3.55E-09 | 3.28E-09 |
| GOTERM_BP_DIRECT | GO:0050832~defense response to fungus | 70 | 11.308562 | 9.80E-11 | 5.55E-09 | 5.13E-09 |
| GOTERM_BP_DIRECT | GO:0009414~response to water deprivation | 68 | 10.98546 | 3.08E-10 | 1.62E-08 | 1.50E-08 |
| GOTERM_BP_DIRECT | GO:0009737~response to abscisic acid | 60 | 9.6930533 | 1.10E-09 | 5.39E-08 | 4.98E-08 |
| GOTERM_BP_DIRECT | GO:0009765~photosynthesis, light harvesting | 10 | 1.6155089 | 1.69E-09 | 7.79E-08 | 7.20E-08 |
| GOTERM_BP_DIRECT | GO:0007165~signal transduction | 51 | 8.2390953 | 1.24E-08 | 5.38E-07 | 4.97E-07 |
| GOTERM_BP_DIRECT | GO:0009617~response to bacterium | 28 | 4.5234249 | 3.00E-08 | 1.23E-06 | 1.13E-06 |
| GOTERM_BP_DIRECT | GO:0009620~response to fungus | 25 | 4.0387722 | 1.34E-07 | 5.20E-06 | 4.80E-06 |
| GOTERM_BP_DIRECT | GO:0006979~response to oxidative stress | 37 | 5.9773829 | 3.71E-07 | 1.37E-05 | 1.26E-05 |
| GOTERM_BP_DIRECT | GO:0009753~response to jasmonic acid | 35 | 5.6542811 | 4.68E-07 | 1.64E-05 | 1.52E-05 |
| GOTERM_BP_DIRECT | GO:0018298~protein-chromophore linkage | 11 | 1.7770598 | 5.12E-07 | 1.65E-05 | 1.52E-05 |
| GOTERM_BP_DIRECT | GO:0071407~cellular response to organic cyclic compound | 14 | 2.2617124 | 5.15E-07 | 1.65E-05 | 1.52E-05 |
| GOTERM_BP_DIRECT | GO:0098542~defense response to other organism | 39 | 6.3004847 | 7.09E-07 | 2.17E-05 | 2.01E-05 |
| GOTERM_BP_DIRECT | GO:0009769~photosynthesis, light harvesting in photosystem II | 5 | 0.8077544 | 5.00E-06 | 1.47E-04 | 1.36E-04 |
| GOTERM_BP_DIRECT | GO:0071310~cellular response to organic substance | 18 | 2.907916 | 5.81E-06 | 1.65E-04 | 1.52E-04 |
| GOTERM_BP_DIRECT | GO:0042542~response to hydrogen peroxide | 11 | 1.7770598 | 6.30E-06 | 1.72E-04 | 1.59E-04 |
| GOTERM_BP_DIRECT | GO:1901615~organic hydroxy compound metabolic process | 8 | 1.2924071 | 1.20E-05 | 3.15E-04 | 2.91E-04 |
| GOTERM_BP_DIRECT | GO:0032787~monocarboxylic acid metabolic process | 16 | 2.5848142 | 2.46E-05 | 6.25E-04 | 5.78E-04 |
| GOTERM_BP_DIRECT | GO:0009651~response to salt stress | 34 | 5.4927302 | 5.85E-05 | 0.0014351 | 0.0013259 |
| GOTERM_BP_DIRECT | GO:0010114~response to red light | 11 | 1.7770598 | 7.38E-05 | 0.0017519 | 0.0016186 |
| GOTERM_BP_DIRECT | GO:0006952~defense response | 38 | 6.1389338 | 1.01E-04 | 0.0023197 | 0.0021432 |
| GOTERM_BP_DIRECT | GO:0071215~cellular response to abscisic acid stimulus | 10 | 1.6155089 | 1.04E-04 | 0.0023197 | 0.0021432 |
| GOTERM_BP_DIRECT | GO:0097305~response to alcohol | 13 | 2.1001616 | 1.55E-04 | 0.0033553 | 0.0031 |
| GOTERM_BP_DIRECT | GO:1990961~drug transmembrane export | 8 | 1.2924071 | 1.96E-04 | 0.0041264 | 0.0038124 |
| GOTERM_BP_DIRECT | GO:0051707~response to other organism | 6 | 0.9693053 | 2.43E-04 | 0.0048286 | 0.0044612 |
| GOTERM_BP_DIRECT | GO:0009645~response to low light intensity stimulus | 6 | 0.9693053 | 2.43E-04 | 0.0048286 | 0.0044612 |
| GOTERM_BP_DIRECT | GO:0009407~toxin catabolic process | 7 | 1.1308562 | 8.28E-04 | 0.0160421 | 0.0148215 |
| GOTERM_BP_DIRECT | GO:1900426~positive regulation of defense response to bacteriun | 6 | 0.9693053 | 9.80E-04 | 0.0180864 | 0.0167103 |
| GOTERM_BP_DIRECT | GO:0002238~response to molecule of fungal origin | 5 | 0.8077544 | 0.001005 | 0.0180864 | 0.0167103 |
| GOTERM_BP_DIRECT | GO:0002229~defense response to oomycetes | 9 | 1.453958 | 0.0010075 | 0.0180864 | 0.0167103 |
| GOTERM_BP_DIRECT | GO:0006749~glutathione metabolic process | 8 | 1.2924071 | 0.0010952 | 0.0191926 | 0.0177323 |
| GOTERM_BP_DIRECT | GO:0010565~regulation of cellular ketone metabolic process | 7 | 1.1308562 | 0.0011628 | 0.0199035 | 0.0183891 |
| GOTERM_BP_DIRECT | GO:0009863~salicylic acid mediated signaling pathway | 8 | 1.2924071 | 0.0011978 | 0.020036 | 0.0185115 |
| GOTERM_BP_DIRECT | GO:0009072~aromatic amino acid family metabolic process | 7 | 1.1308562 | 0.0017592 | 0.0287721 | 0.0265829 |
| GOTERM_BP_DIRECT | GO:0006468~protein phosphorylation | 37 | 5.9773829 | 0.0019991 | 0.0319851 | 0.0295514 |
| GOTERM_BP_DIRECT | GO:0009626~plant-type hypersensitive response | 9 | 1.453958 | 0.002228 | 0.0348896 | 0.032235 |
| GOTERM_BP_DIRECT | GO:0009409~response to cold | 27 | 4.361874 | 0.0024205 | 0.037114 | 0.0342901 |
| GOTERM_BP_DIRECT | GO:0043436~oxoacid metabolic process | 18 | 2.907916 | 0.0029118 | 0.0437361 | 0.0404083 |

| | | | | | | |
|---|---|---|---|---|---|---|
| GOTERM_BP_DIRECT | GO:0009269~response to desiccation | 5 | 0.8077544 | 0.0033881 | 0.0495366 | 0.0457676 |
| GOTERM_BP_DIRECT | GO:1903428~positive regulation of reactive oxygen species biosy | 3 | 0.4846527 | 0.0034326 | 0.0495366 | 0.0457676 |
| GOTERM_BP_DIRECT | GO:0006970~response to osmotic stress | 21 | 3.3925687 | 0.0036575 | 0.0517679 | 0.047829 |
| GOTERM_BP_DIRECT | GO:0032870~cellular response to hormone stimulus | 11 | 1.7770598 | 0.0037705 | 0.0523595 | 0.0483756 |
| GOTERM_BP_DIRECT | GO:0071446~cellular response to salicylic acid stimulus | 6 | 0.9693053 | 0.0045886 | 0.0625407 | 0.0577822 |
| GOTERM_BP_DIRECT | GO:0009695~jasmonic acid biosynthetic process | 5 | 0.8077544 | 0.0050823 | 0.0680098 | 0.0628352 |
| GOTERM_BP_DIRECT | GO:0001666~response to hypoxia | 10 | 1.6155089 | 0.00558 | 0.0733369 | 0.0677569 |
| GOTERM_BP_DIRECT | GO:0031408~oxylipin biosynthetic process | 5 | 0.8077544 | 0.0057531 | 0.074286 | 0.0686338 |
| GOTERM_BP_DIRECT | GO:0016998~cell wall macromolecule catabolic process | 4 | 0.6462036 | 0.0063358 | 0.0803996 | 0.0742823 |
| GOTERM_BP_DIRECT | GO:0009812~flavonoid metabolic process | 7 | 1.1308562 | 0.0066344 | 0.0827618 | 0.0764647 |
| GOTERM_BP_DIRECT | GO:0010224~response to UV-B | 7 | 1.1308562 | 0.007111 | 0.0855642 | 0.0790538 |
| GOTERM_BP_DIRECT | GO:0071396~cellular response to lipid | 17 | 2.7463651 | 0.0071472 | 0.0855642 | 0.0790538 |
| GOTERM_BP_DIRECT | GO:0009644~response to high light intensity | 6 | 0.9693053 | 0.0072079 | 0.0855642 | 0.0790538 |
| GOTERM_BP_DIRECT | GO:0009416~response to light stimulus | 43 | 6.9466882 | 0.0078788 | 0.0920445 | 0.0850411 |
| GOTERM_BP_DIRECT | GO:1901362~organic cyclic compound biosynthetic process | 22 | 3.5541195 | 0.0098478 | 0.1132503 | 0.1046334 |
| GOTERM_BP_DIRECT | GO:0033993~response to lipid | 29 | 4.6849758 | 0.0110053 | 0.1246139 | 0.1151324 |
| GOTERM_BP_DIRECT | GO:0019438~aromatic compound biosynthetic process | 20 | 3.2310178 | 0.011775 | 0.1313086 | 0.1213177 |
| GOTERM_BP_DIRECT | GO:0098771~inorganic ion homeostasis | 7 | 1.1308562 | 0.0125705 | 0.1361791 | 0.1258176 |
| GOTERM_BP_DIRECT | GO:0005975~carbohydrate metabolic process | 19 | 3.0694669 | 0.0125818 | 0.1361791 | 0.1258176 |
| GOTERM_BP_DIRECT | GO:0009867~jasmonic acid mediated signaling pathway | 10 | 1.6155089 | 0.0154734 | 0.1644395 | 0.1519278 |
| GOTERM_BP_DIRECT | GO:0009636~response to toxic substance | 4 | 0.6462036 | 0.0156429 | 0.1644395 | 0.1519278 |
| GOTERM_BP_DIRECT | GO:0009615~response to virus | 5 | 0.8077544 | 0.0158631 | 0.1644395 | 0.1519278 |
| GOTERM_BP_DIRECT | GO:0090333~regulation of stomatal closure | 5 | 0.8077544 | 0.0172486 | 0.1763186 | 0.1629031 |
| GOTERM_BP_DIRECT | GO:0030104~water homeostasis | 3 | 0.4846527 | 0.0189963 | 0.1915248 | 0.1769523 |
| GOTERM_BP_DIRECT | GO:0006032~chitin catabolic process | 4 | 0.6462036 | 0.0198374 | 0.1973013 | 0.1822892 |
| GOTERM_BP_DIRECT | GO:0014070~response to organic cyclic compound | 16 | 2.5848142 | 0.0221922 | 0.2177799 | 0.2012097 |
| GOTERM_BP_DIRECT | GO:2000068~regulation of defense response to insect | 3 | 0.4846527 | 0.0233671 | 0.2262916 | 0.2090737 |
| GOTERM_BP_DIRECT | GO:0016051~carbohydrate biosynthetic process | 4 | 0.6462036 | 0.0299017 | 0.2858133 | 0.2640666 |
| GOTERM_BP_DIRECT | GO:0010093~specification of floral organ identity | 3 | 0.4846527 | 0.0331916 | 0.3131923 | 0.2893624 |
| GOTERM_BP_DIRECT | GO:0009058~biosynthetic process | 6 | 0.9693053 | 0.0342501 | 0.3190894 | 0.2948109 |
| GOTERM_BP_DIRECT | GO:0009755~hormone-mediated signaling pathway | 14 | 2.2617124 | 0.0354749 | 0.3263694 | 0.3015369 |
| GOTERM_BP_DIRECT | GO:0019748~secondary metabolic process | 11 | 1.7770598 | 0.0374517 | 0.3365235 | 0.3109185 |
| GOTERM_BP_DIRECT | GO:0010218~response to far red light | 5 | 0.8077544 | 0.0374931 | 0.3365235 | 0.3109185 |
| GOTERM_BP_DIRECT | GO:0010112~regulation of systemic acquired resistance | 3 | 0.4846527 | 0.0386053 | 0.3423313 | 0.3162844 |
| GOTERM_BP_DIRECT | GO:1900409~positive regulation of cellular response to oxidative s | 2 | 0.3231018 | 0.0480728 | 0.4114141 | 0.3801109 |
| GOTERM_BP_DIRECT | GO:0046256~2,4,6-trinitrotoluene catabolic process | 2 | 0.3231018 | 0.0480728 | 0.4114141 | 0.3801109 |
| GOTERM_BP_DIRECT | GO:0010249~auxin conjugate metabolic process | 2 | 0.3231018 | 0.0480728 | 0.4114141 | 0.3801109 |
| GOTERM_BP_DIRECT | GO:0031349~positive regulation of defense response | 3 | 0.4846527 | 0.0503412 | 0.4258746 | 0.3934711 |
| GOTERM_BP_DIRECT | GO:0006629~lipid metabolic process | 11 | 1.7770598 | 0.0534242 | 0.4468206 | 0.4128233 |
| GOTERM_BP_DIRECT | GO:0031668~cellular response to extracellular stimulus | 3 | 0.4846527 | 0.0631705 | 0.5165943 | 0.4772882 |
| GOTERM_BP_DIRECT | GO:0046283~anthocyanin-containing compound metabolic proce | 3 | 0.4846527 | 0.0631705 | 0.5165943 | 0.4772882 |
| GOTERM_BP_DIRECT | GO:0033014~tetrapyrrole biosynthetic process | 4 | 0.6462036 | 0.0687227 | 0.5558228 | 0.5135319 |

**Table S3. GO biological Processes enrichment for 16:8 and CLL-common DEGs.**

| Category | Term | Count | % | PValue | Benjamini | FDR |
|---|---|---|---|---|---|---|
| GOTERM_BP_DIRECT | GO:0009733~response to auxin | 107 | 3.372203 | 2.04E-20 | 4.00E-17 | 3.86E-17 |
| GOTERM_BP_DIRECT | GO:0009416~response to light stimulus | 229 | 7.217145 | 1.81E-13 | 1.77E-10 | 1.71E-10 |
| GOTERM_BP_DIRECT | GO:0010374~stomatal complex development | 24 | 0.756382 | 9.94E-12 | 6.48E-09 | 6.26E-09 |
| GOTERM_BP_DIRECT | GO:0015979~photosynthesis | 48 | 1.512764 | 4.77E-10 | 2.33E-07 | 2.25E-07 |
| GOTERM_BP_DIRECT | GO:0033993~response to lipid | 146 | 4.601324 | 7.31E-10 | 2.86E-07 | 2.76E-07 |
| GOTERM_BP_DIRECT | GO:0019761~glucosinolate biosynthetic process | 36 | 1.134573 | 1.18E-09 | 3.83E-07 | 3.70E-07 |
| GOTERM_BP_DIRECT | GO:0009642~response to light intensity | 63 | 1.985503 | 1.60E-09 | 4.46E-07 | 4.31E-07 |
| GOTERM_BP_DIRECT | GO:0009611~response to wounding | 172 | 5.420737 | 3.50E-09 | 8.56E-07 | 8.27E-07 |
| GOTERM_BP_DIRECT | GO:0009734~auxin-activated signaling pathway | 60 | 1.890955 | 1.43E-08 | 2.94E-06 | 2.84E-06 |
| GOTERM_BP_DIRECT | GO:0009753~response to jasmonic acid | 112 | 3.529783 | 1.50E-08 | 2.94E-06 | 2.84E-06 |
| GOTERM_BP_DIRECT | GO:0007018~microtubule-based movement | 26 | 0.819414 | 2.44E-07 | 4.35E-05 | 4.20E-05 |
| GOTERM_BP_DIRECT | GO:1903046~meiotic cell cycle process | 40 | 1.260637 | 3.70E-07 | 6.04E-05 | 5.83E-05 |
| GOTERM_BP_DIRECT | GO:0051301~cell division | 103 | 3.246139 | 4.52E-07 | 6.80E-05 | 6.56E-05 |
| GOTERM_BP_DIRECT | GO:0009414~response to water deprivation | 206 | 6.492279 | 5.27E-07 | 7.36E-05 | 7.11E-05 |
| GOTERM_BP_DIRECT | GO:0032787~monocarboxylic acid metabolic process | 46 | 1.449732 | 6.31E-07 | 8.23E-05 | 7.95E-05 |
| GOTERM_BP_DIRECT | GO:0044271~cellular nitrogen compound biosynthetic | 81 | 2.552789 | 1.70E-06 | 2.08E-04 | 2.01E-04 |
| GOTERM_BP_DIRECT | GO:0019438~aromatic compound biosynthetic proces | 86 | 2.710369 | 9.89E-06 | 0.001138 | 0.001099 |
| GOTERM_BP_DIRECT | GO:0006979~response to oxidative stress | 107 | 3.372203 | 1.56E-05 | 0.001692 | 0.001634 |
| GOTERM_BP_DIRECT | GO:0042343~indole glucosinolate metabolic process | 13 | 0.409707 | 1.98E-05 | 0.002035 | 0.001965 |
| GOTERM_BP_DIRECT | GO:0009888~tissue development | 132 | 4.160101 | 2.17E-05 | 0.002098 | 0.002026 |
| GOTERM_BP_DIRECT | GO:0051321~meiotic cell cycle | 22 | 0.69335 | 2.25E-05 | 0.002098 | 0.002026 |
| GOTERM_BP_DIRECT | GO:0009751~response to salicylic acid | 63 | 1.985503 | 2.52E-05 | 0.002241 | 0.002164 |
| GOTERM_BP_DIRECT | GO:0010143~cutin biosynthetic process | 12 | 0.378191 | 2.91E-05 | 0.002476 | 0.002391 |
| GOTERM_BP_DIRECT | GO:1901362~organic cyclic compound biosynthetic pr | 92 | 2.899464 | 3.16E-05 | 0.002574 | 0.002486 |
| GOTERM_BP_DIRECT | GO:0009059~macromolecule biosynthetic process | 163 | 5.137094 | 3.89E-05 | 0.003048 | 0.002944 |
| GOTERM_BP_DIRECT | GO:0015995~chlorophyll biosynthetic process | 23 | 0.724866 | 5.91E-05 | 0.004446 | 0.004294 |
| GOTERM_BP_DIRECT | GO:0044772~mitotic cell cycle phase transition | 14 | 0.441223 | 6.93E-05 | 0.004794 | 0.00463 |
| GOTERM_BP_DIRECT | GO:0019253~reductive pentose-phosphate cycle | 11 | 0.346675 | 7.10E-05 | 0.004794 | 0.00463 |
| GOTERM_BP_DIRECT | GO:0009741~response to brassinosteroid | 19 | 0.598802 | 7.33E-05 | 0.004794 | 0.00463 |
| GOTERM_BP_DIRECT | GO:0007076~mitotic chromosome condensation | 7 | 0.220611 | 7.40E-05 | 0.004794 | 0.00463 |
| GOTERM_BP_DIRECT | GO:0010817~regulation of hormone levels | 12 | 0.378191 | 7.59E-05 | 0.004794 | 0.00463 |
| GOTERM_BP_DIRECT | GO:0040008~regulation of growth | 36 | 1.134573 | 8.03E-05 | 0.004908 | 0.00474 |
| GOTERM_BP_DIRECT | GO:0009266~response to temperature stimulus | 68 | 2.143082 | 1.39E-04 | 0.008241 | 0.007959 |
| GOTERM_BP_DIRECT | GO:1901673~regulation of mitotic spindle assembly | 6 | 0.189095 | 1.46E-04 | 0.008295 | 0.008011 |
| GOTERM_BP_DIRECT | GO:0008610~lipid biosynthetic process | 55 | 1.733375 | 1.49E-04 | 0.008295 | 0.008011 |
| GOTERM_BP_DIRECT | GO:0043436~oxoacid metabolic process | 64 | 2.017019 | 1.53E-04 | 0.008295 | 0.008011 |
| GOTERM_BP_DIRECT | GO:0009773~photosynthetic electron transport in pho | 10 | 0.315159 | 1.74E-04 | 0.009187 | 0.008873 |
| GOTERM_BP_DIRECT | GO:0040007~growth | 42 | 1.323668 | 1.84E-04 | 0.009488 | 0.009163 |
| GOTERM_BP_DIRECT | GO:0009737~response to abscisic acid | 165 | 5.200126 | 1.96E-04 | 0.009821 | 0.009485 |
| GOTERM_BP_DIRECT | GO:0009639~response to red or far red light | 41 | 1.292153 | 2.10E-04 | 0.010266 | 0.009914 |
| GOTERM_BP_DIRECT | GO:0007623~circadian rhythm | 31 | 0.976993 | 2.21E-04 | 0.010536 | 0.010176 |
| GOTERM_BP_DIRECT | GO:0022607~cellular component assembly | 24 | 0.756382 | 2.31E-04 | 0.010754 | 0.010386 |
| GOTERM_BP_DIRECT | GO:0009735~response to cytokinin | 20 | 0.630318 | 2.41E-04 | 0.010988 | 0.010612 |
| GOTERM_BP_DIRECT | GO:0071456~cellular response to hypoxia | 50 | 1.575796 | 2.63E-04 | 0.011719 | 0.011318 |
| GOTERM_BP_DIRECT | GO:0000079~regulation of cyclin-dependent protein s | 16 | 0.504255 | 3.13E-04 | 0.013625 | 0.013159 |
| GOTERM_BP_DIRECT | GO:0002213~defense response to insect | 21 | 0.661834 | 4.21E-04 | 0.017782 | 0.017174 |
| GOTERM_BP_DIRECT | GO:0010052~guard cell differentiation | 9 | 0.283643 | 4.27E-04 | 0.017782 | 0.017174 |
| GOTERM_BP_DIRECT | GO:0007094~mitotic spindle assembly checkpoint | 7 | 0.220611 | 4.46E-04 | 0.018183 | 0.017561 |
| GOTERM_BP_DIRECT | GO:0080028~nitrile biosynthetic process | 6 | 0.189095 | 4.58E-04 | 0.018312 | 0.017685 |
| GOTERM_BP_DIRECT | GO:0055072~iron ion homeostasis | 15 | 0.472739 | 5.10E-04 | 0.019963 | 0.01928 |
| GOTERM_BP_DIRECT | GO:0048366~leaf development | 41 | 1.292153 | 5.82E-04 | 0.022316 | 0.021552 |
| GOTERM_BP_DIRECT | GO:1901566~organonitrogen compound biosynthetic | 53 | 1.670344 | 6.03E-04 | 0.022543 | 0.021771 |
| GOTERM_BP_DIRECT | GO:0006631~fatty acid metabolic process | 26 | 0.819414 | 6.11E-04 | 0.022543 | 0.021771 |
| GOTERM_BP_DIRECT | GO:0009736~cytokinin-activated signaling pathway | 24 | 0.756382 | 7.62E-04 | 0.027611 | 0.026665 |
| GOTERM_BP_DIRECT | GO:0018130~heterocycle biosynthetic process | 40 | 1.260637 | 8.22E-04 | 0.029266 | 0.028264 |

| | | | | | | |
|---|---|---|---|---|---|---|
| [illegible] | [illegible] | | | | | |
| GOTERM_BP_DIRECT | GO:0019684~photosynthesis, light reaction | 15 | 0.472739 | 8.41E-04 | 0.029387 | 0.028381 |
| GOTERM_BP_DIRECT | GO:0006355~regulation of transcription, DNA-templat | 274 | 8.635361 | 9.42E-04 | 0.032358 | 0.03125 |
| GOTERM_BP_DIRECT | GO:0032465~regulation of cytokinesis | 5 | 0.15758 | 0.001001 | 0.033762 | 0.032606 |
| GOTERM_BP_DIRECT | GO:0009625~response to insect | 14 | 0.441223 | 0.001072 | 0.035549 | 0.034332 |
| GOTERM_BP_DIRECT | GO:0042742~defense response to bacterium | 167 | 5.263158 | 0.001144 | 0.037322 | 0.036044 |
| GOTERM_BP_DIRECT | GO:0009556~microsporogenesis | 11 | 0.346675 | 0.001178 | 0.037792 | 0.036498 |
| GOTERM_BP_DIRECT | GO:0060586~multicellular organismal iron ion homeos | 11 | 0.346675 | 0.001603 | 0.050604 | 0.048872 |
| GOTERM_BP_DIRECT | GO:0000910~cytokinesis | 21 | 0.661834 | 0.001701 | 0.052846 | 0.051037 |
| GOTERM_BP_DIRECT | GO:0007049~cell cycle | 47 | 1.481248 | 0.001745 | 0.05337 | 0.051542 |
| GOTERM_BP_DIRECT | GO:0009409~response to cold | 96 | 3.025528 | 0.002052 | 0.061793 | 0.059677 |
| GOTERM_BP_DIRECT | GO:0071446~cellular response to salicylic acid stimulu | 14 | 0.441223 | 0.00216 | 0.064034 | 0.061842 |
| GOTERM_BP_DIRECT | GO:0009641~shade avoidance | 8 | 0.252127 | 0.002557 | 0.074699 | 0.072142 |
| GOTERM_BP_DIRECT | GO:0043687~post-translational protein modification | 5 | 0.15758 | 0.00271 | 0.07672 | 0.074093 |
| GOTERM_BP_DIRECT | GO:0014070~response to organic cyclic compound | 61 | 1.922471 | 0.002734 | 0.07672 | 0.074093 |
| GOTERM_BP_DIRECT | GO:0010114~response to red light | 22 | 0.69335 | 0.002744 | 0.07672 | 0.074093 |
| GOTERM_BP_DIRECT | GO:0031347~regulation of defense response | 103 | 3.246139 | 0.002948 | 0.081267 | 0.078484 |
| GOTERM_BP_DIRECT | GO:0002237~response to molecule of bacterial origin | 32 | 1.008509 | 0.003008 | 0.081765 | 0.078966 |
| GOTERM_BP_DIRECT | GO:0009624~response to nematode | 21 | 0.661834 | 0.003108 | 0.083317 | 0.080465 |
| GOTERM_BP_DIRECT | GO:0009767~photosynthetic electron transport chain | 9 | 0.283643 | 0.003203 | 0.083566 | 0.080705 |
| GOTERM_BP_DIRECT | GO:0010345~suberin biosynthetic process | 9 | 0.283643 | 0.003203 | 0.083566 | 0.080705 |
| GOTERM_BP_DIRECT | GO:0009651~response to salt stress | 103 | 3.246139 | 0.003903 | 0.099688 | 0.096275 |
| GOTERM_BP_DIRECT | GO:0010103~stomatal complex morphogenesis | 6 | 0.189095 | 0.004003 | 0.099688 | 0.096275 |
| GOTERM_BP_DIRECT | GO:0010389~regulation of G2/M transition of mitotic | 6 | 0.189095 | 0.004003 | 0.099688 | 0.096275 |
| GOTERM_BP_DIRECT | GO:0009813~flavonoid biosynthetic process | 14 | 0.441223 | 0.004024 | 0.099688 | 0.096275 |
| GOTERM_BP_DIRECT | GO:0009635~response to herbicide | 7 | 0.220611 | 0.004129 | 0.100998 | 0.09754 |
| GOTERM_BP_DIRECT | GO:0010120~camalexin biosynthetic process | 8 | 0.252127 | 0.005316 | 0.128434 | 0.124037 |
| GOTERM_BP_DIRECT | GO:0009645~response to low light intensity stimulus | 9 | 0.283643 | 0.005892 | 0.138933 | 0.134177 |
| GOTERM_BP_DIRECT | GO:0009718~anthocyanin-containing compound bios | 9 | 0.283643 | 0.005892 | 0.138933 | 0.134177 |
| GOTERM_BP_DIRECT | GO:0099402~plant organ development | 13 | 0.409707 | 0.006424 | 0.149669 | 0.144545 |
| GOTERM_BP_DIRECT | GO:0010444~guard mother cell differentiation | 6 | 0.189095 | 0.006606 | 0.152094 | 0.146886 |
| GOTERM_BP_DIRECT | GO:0009826~unidimensional cell growth | 41 | 1.292153 | 0.006846 | 0.154679 | 0.149383 |
| GOTERM_BP_DIRECT | GO:0009755~hormone-mediated signaling pathway | 53 | 1.670344 | 0.006938 | 0.154679 | 0.149383 |
| GOTERM_BP_DIRECT | GO:0015996~chlorophyll catabolic process | 12 | 0.378191 | 0.006955 | 0.154679 | 0.149383 |
| GOTERM_BP_DIRECT | GO:0010035~response to inorganic substance | 84 | 2.647337 | 0.00725 | 0.15942 | 0.153962 |
| GOTERM_BP_DIRECT | GO:0003002~regionalization | 15 | 0.472739 | 0.007443 | 0.161848 | 0.156307 |
| GOTERM_BP_DIRECT | GO:0044255~cellular lipid metabolic process | 44 | 1.3867 | 0.007786 | 0.167444 | 0.161712 |
| GOTERM_BP_DIRECT | GO:0048869~cellular developmental process | 20 | 0.630318 | 0.007956 | 0.169245 | 0.16345 |
| GOTERM_BP_DIRECT | GO:0010150~leaf senescence | 50 | 1.575796 | 0.008838 | 0.184779 | 0.178453 |
| GOTERM_BP_DIRECT | GO:0051026~chiasma assembly | 7 | 0.220611 | 0.008875 | 0.184779 | 0.178453 |
| GOTERM_BP_DIRECT | GO:0044550~secondary metabolite biosynthetic proce | 29 | 0.913962 | 0.009094 | 0.186806 | 0.180411 |
| GOTERM_BP_DIRECT | GO:0009851~auxin biosynthetic process | 11 | 0.346675 | 0.009164 | 0.186806 | 0.180411 |
| GOTERM_BP_DIRECT | GO:0042435~indole-containing compound biosynthet | 15 | 0.472739 | 0.010196 | 0.1957 | 0.189 |
| GOTERM_BP_DIRECT | GO:0043933~macromolecular complex subunit organi: | 6 | 0.189095 | 0.010198 | 0.1957 | 0.189 |
| GOTERM_BP_DIRECT | GO:0010106~cellular response to iron ion starvation | 5 | 0.15758 | 0.010326 | 0.1957 | 0.189 |
| GOTERM_BP_DIRECT | GO:0010258~NADH dehydrogenase complex (plastoqu | 5 | 0.15758 | 0.010326 | 0.1957 | 0.189 |
| GOTERM_BP_DIRECT | GO:0000082~G1/S transition of mitotic cell cycle | 5 | 0.15758 | 0.010326 | 0.1957 | 0.189 |
| GOTERM_BP_DIRECT | GO:0042549~photosystem II stabilization | 5 | 0.15758 | 0.010326 | 0.1957 | 0.189 |
| GOTERM_BP_DIRECT | GO:0048232~male gamete generation | 5 | 0.15758 | 0.010326 | 0.1957 | 0.189 |
| GOTERM_BP_DIRECT | GO:0044248~cellular catabolic process | 70 | 2.206114 | 0.0104 | 0.1957 | 0.189 |
| GOTERM_BP_DIRECT | GO:0000911~cytokinesis by cell plate formation | 13 | 0.409707 | 0.011021 | 0.205408 | 0.198375 |
| GOTERM_BP_DIRECT | GO:0080090~regulation of primary metabolic process | 54 | 1.701859 | 0.011419 | 0.210828 | 0.20361 |
| GOTERM_BP_DIRECT | GO:0000160~phosphorelay signal transduction systen | 12 | 0.378191 | 0.012287 | 0.221224 | 0.21365 |
| GOTERM_BP_DIRECT | GO:1990641~response to iron ion starvation | 7 | 0.220611 | 0.012322 | 0.221224 | 0.21365 |
| GOTERM_BP_DIRECT | GO:0051259~protein oligomerization | 7 | 0.220611 | 0.012322 | 0.221224 | 0.21365 |
| GOTERM_BP_DIRECT | GO:0006833~water transport | 8 | 0.252127 | 0.012956 | 0.230501 | 0.222609 |

**Table S4. GO biological Processes enrichment for 16:8-exclusive DEGs.**

| Category | Term | Count | % | PValue | Benjamini | FDR |
|---|---|---|---|---|---|---|
| GOTERM_BF | GO:0009751~response to salicylic acid | 24 | 4.75247525 | 4.13E-08 | 1.56E-05 | 1.53E-05 |
| GOTERM_BF | GO:0009611~response to wounding | 45 | 8.91089109 | 4.81E-08 | 1.56E-05 | 1.53E-05 |
| GOTERM_BF | GO:0042742~defense response to bacterium | 50 | 9.9009901 | 7.02E-08 | 1.56E-05 | 1.53E-05 |
| GOTERM_BF | GO:0060586~multicellular organismal iron ion homeostasis | 8 | 1.58415842 | 1.21E-06 | 2.01E-04 | 1.98E-04 |
| GOTERM_BF | GO:0055072~iron ion homeostasis | 9 | 1.78217822 | 2.08E-06 | 2.78E-04 | 2.73E-04 |
| GOTERM_BF | GO:0010106~cellular response to iron ion starvation | 5 | 0.99009901 | 9.90E-06 | 0.00110065 | 0.0010825 |
| GOTERM_BF | GO:0050832~defense response to fungus | 48 | 9.5049505 | 1.35E-05 | 0.00113673 | 0.00111798 |
| GOTERM_BF | GO:0031347~regulation of defense response | 31 | 6.13861386 | 1.36E-05 | 0.00113673 | 0.00111798 |
| GOTERM_BF | GO:1901701~cellular response to oxygen-containing compound | 26 | 5.14851485 | 1.97E-05 | 0.00146019 | 0.00143611 |
| GOTERM_BF | GO:0002237~response to molecule of bacterial origin | 12 | 2.37623762 | 2.41E-04 | 0.01605303 | 0.01578829 |
| GOTERM_BF | GO:0009753~response to jasmonic acid | 24 | 4.75247525 | 4.99E-04 | 0.03028594 | 0.02978647 |
| GOTERM_BF | GO:0009414~response to water deprivation | 41 | 8.11881188 | 0.00110096 | 0.06119512 | 0.06018591 |
| GOTERM_BF | GO:0010039~response to iron ion | 5 | 0.99009901 | 0.00241801 | 0.11965436 | 0.11768105 |
| GOTERM_BF | GO:0009813~flavonoid biosynthetic process | 6 | 1.18811881 | 0.00251149 | 0.11965436 | 0.11768105 |
| GOTERM_BF | GO:0009733~response to auxin | 16 | 3.16831683 | 0.00373047 | 0.16588139 | 0.16314571 |
| GOTERM_BF | GO:0010150~leaf senescence | 14 | 2.77227723 | 0.00474636 | 0.1978638 | 0.19460068 |
| GOTERM_BF | GO:0033993~response to lipid | 26 | 5.14851485 | 0.00518439 | 0.20341124 | 0.20005663 |
| GOTERM_BF | GO:0009737~response to abscisic acid | 33 | 6.53465347 | 0.0083332 | 0.28050377 | 0.27587777 |
| GOTERM_BF | GO:0010035~response to inorganic substance | 20 | 3.96039604 | 0.00850578 | 0.28050377 | 0.27587777 |
| GOTERM_BF | GO:0019760~glucosinolate metabolic process | 8 | 1.58415842 | 0.00907535 | 0.28050377 | 0.27587777 |
| GOTERM_BF | GO:0009625~response to insect | 5 | 0.99009901 | 0.00931886 | 0.28050377 | 0.27587777 |
| GOTERM_BF | GO:0009416~response to light stimulus | 36 | 7.12871287 | 0.00998656 | 0.28050377 | 0.27587777 |
| GOTERM_BF | GO:0010258~NADH dehydrogenase complex (plastoquinone) assem | 3 | 0.59405941 | 0.01008625 | 0.28050377 | 0.27587777 |
| GOTERM_BF | GO:0015979~photosynthesis | 9 | 1.78217822 | 0.01009309 | 0.28050377 | 0.27587777 |
| GOTERM_BF | GO:0071446~cellular response to salicylic acid stimulus | 5 | 0.99009901 | 0.01184529 | 0.31555671 | 0.31035263 |
| GOTERM_BF | GO:0006979~response to oxidative stress | 21 | 4.15841584 | 0.0126191 | 0.31555671 | 0.31035263 |
| GOTERM_BF | GO:0006879~cellular iron ion homeostasis | 4 | 0.79207921 | 0.01277366 | 0.31555671 | 0.31035263 |
| GOTERM_BF | GO:0044272~sulfur compound biosynthetic process | 6 | 1.18811881 | 0.01559276 | 0.35863354 | 0.35271905 |
| GOTERM_BF | GO:0002213~defense response to insect | 6 | 1.18811881 | 0.01559276 | 0.35863354 | 0.35271905 |
| GOTERM_BF | GO:0009755~hormone-mediated signaling pathway | 13 | 2.57425743 | 0.01780144 | 0.38681177 | 0.38043257 |
| GOTERM_BF | GO:0045087~innate immune response | 8 | 1.58415842 | 0.01797776 | 0.38681177 | 0.38043257 |
| GOTERM_BF | GO:0043455~regulation of secondary metabolic process | 4 | 0.79207921 | 0.02097359 | 0.43716822 | 0.42995855 |
| GOTERM_BF | GO:0071456~cellular response to hypoxia | 11 | 2.17821782 | 0.02306337 | 0.46615972 | 0.45847193 |
| GOTERM_BF | GO:0009409~response to cold | 20 | 3.96039604 | 0.02417897 | 0.47433452 | 0.46651191 |
| GOTERM_BF | GO:0010584~pollen exine formation | 4 | 0.79207921 | 0.0269922 | 0.5143943 | 0.50591104 |
| GOTERM_BF | GO:0071396~cellular response to lipid | 13 | 2.57425743 | 0.03082601 | 0.54602637 | 0.53702144 |
| GOTERM_BF | GO:0010114~response to red light | 6 | 1.18811881 | 0.03441907 | 0.54602637 | 0.53702144 |
| GOTERM_BF | GO:0006880~intracellular sequestering of iron ion | 3 | 0.59405941 | 0.03453107 | 0.54602637 | 0.53702144 |
| GOTERM_BF | GO:0006949~syncytium formation | 3 | 0.59405941 | 0.03453107 | 0.54602637 | 0.53702144 |
| GOTERM_BF | GO:0006826~iron ion transport | 3 | 0.59405941 | 0.03453107 | 0.54602637 | 0.53702144 |

**Table S5. GO Biological Processes enrichment for 509 DEGs in Fig. 8.**
